# Distinct evolutionary trajectories of SARS-CoV-2 interacting proteins in bats and primates identify important host determinants of COVID-19

**DOI:** 10.1101/2022.04.07.487460

**Authors:** Marie Cariou, Léa Picard, Laurent Guéguen, Stéphanie Jacquet, Andrea Cimarelli, Oliver I Fregoso, Antoine Molaro, Vincent Navratil, Lucie Etienne

## Abstract

The COVID-19 pandemic is caused by SARS-CoV-2, a novel coronavirus that spilled from the bat reservoir. Despite numerous clinical trials and vaccines, the burden remains immense, and the host determinants of SARS-CoV-2 susceptibility and COVID-19 severity remain largely unknown. Signatures of positive selection detected by comparative functional-genetic analyses in primate and bat genomes can uncover important and specific adaptations that occurred at virus-host interfaces. Here, we performed high-throughput evolutionary analyses of 334 SARS-CoV-2 interacting proteins to identify SARS-CoV adaptive loci and uncover functional differences between modern humans, primates and bats. Using DGINN (Detection of Genetic INNovation), we identified 38 bat and 81 primate proteins with marks of positive selection. Seventeen genes, including the ACE2 receptor, present adaptive marks in both mammalian orders, suggesting common virus-host interfaces and past epidemics of coronaviruses shaping their genomes. Yet, 84 genes presented distinct adaptations in bats and primates. Notably, residues involved in ubiquitination and phosphorylation of the inflammatory RIPK1 have rapidly evolved in bats but not primates, suggesting different inflammation regulation versus humans. Furthermore, we discovered residues with typical virus-host arms-race marks in primates, such as in the entry factor TMPRSS2 or the autophagy adaptor FYCO1, pointing to host-specific *in vivo* important interfaces that may be drug targets. Finally, we found that FYCO1 sites under adaptation in primates are those associated with severe COVID-19, supporting their importance in pathogenesis and replication. Overall, we identified functional adaptations involved in SARS- CoV-2 infection in bats and primates, critically enlightening modern genetic determinants of virus susceptibility and severity.

**Key findings:**

- Evolutionary history of 334 SARS-CoV-2 interacting proteins (VIPs) in bats and primates identifying how the past has shaped modern viral reservoirs and humans – results publicly-available in an online resource.
- Identification of 81 primate and 38 bat VIPs with signatures of adaptive evolution. The common ones among species delineate a core adaptive interactome, while the ones displaying distinct evolutionary trajectories enlighten host lineage-specific determinants.
- Evidence of primate specific adaptation of the entry factor TMPRSS2 pointing to its host-specific *in vivo* importance and predicting molecular interfaces.
- FYCO1 sites associated with severe COVID-19 in human (GWAS) display hallmarks of ancient adaptive evolution in primates, highlighting its importance in SARS-CoV-2 replication or pathogenesis and differences with the bat reservoir.
- Identification of adaptive evolution in the bat’s multifunctional RIPK1 at residues that may differentially regulate inflammation.

## Introduction

The current COVID-19 pandemic already led to over six million human deaths (WHO April 2022). The causative agent is a novel severe acute respiratory syndrome coronavirus strain, SARS-CoV-2, that originated from viral cross-species transmission from the bat reservoir, directly or through an intermediate host, to human (Temmam et al., 2022). Bats naturally hosts some of the most high-profile zoonotic viruses, including SARS-CoVs, without apparent symptoms (Wang and Anderson, 2019). Despite scores of clinical trials and effective vaccines, the burden from COVID-19 remains immense in humans, and the determinants of SARS-CoV-2 susceptibility and COVID-19 severity remain largely unknown. A powerful way to identify these factors is to use comparative functional genomics to map host-virus interfaces that underly infections in the bat reservoir and the primate host (Christie et al., 2021).

During infection, viruses interact with many host proteins, or viral-interacting proteins (VIPs). While some VIPs are usurped for viral replication away from their “normal” host functions, some are specifically targeting the virus as part of the host antiviral immune defense. Since the emergence of SARS-CoV-2, VIPs have been identified in hundreds of screens using *in vitro* approaches, such as CRISPR/KO screens, cDNA library screens or mass-spectrometry analyses (Gordon et al., 2020; Parkinson et al., 2020). However, the *in vivo* importance of the identified SARS-CoV-2 VIPs remains largely unknown.

From an evolutionary standpoint, the fitness cost imposed by pathogenetic viruses triggers strong selective pressures on VIPs, such that those VIPs able to prevent, or better counteract, viral infection will quickly become fixed in host populations. In turn, host adaptations push viral proteins into recurring counter-adaptations cycles creating stereotypical “virus-host molecular arms-races”. These arms-races are witnessed by signatures of accelerated rates of evolution, or positive selection, over functionally important residues and domains in VIPs (Daugherty and Malik, 2012; Duggal and Emerman, 2012; Enard et al., 2016). Thus, when combined with functional data, identifying the VIPs with signatures of positive selection is a powerful way to discover virus-host interfaces (e.g. (Boys et al., 2020; Fregoso et al., 2013; Sawyer et al., 2005)).

When studies of adaptive signatures in host genes are combined with human clinical studies or genome-wide association studies (GWAS), they are powerful to uncover the importance of gene evolution and variants in disease severity (e.g. (Wickenhagen et al., 2021; Xie et al., 2018)). Interestingly, several genetic loci associated with COVID-19 severity and susceptibility in humans, such as OAS1 (2’-5’-Oligoadenylate Synthetase 1) or those from the interferon signaling pathway (Bastard et al., 2020; Crow and Stetson, 2021; Schoggins, 2021; The Severe Covid-19 GWAS Group, 2020; Wickenhagen et al., 2021; Zeberg and Pääbo, 2020; Zhang et al., 2020), bear hallmarks of such adaptive arms-races. Furthermore, dozens of VIPs that bear marks of adaptive evolution in the human lineage from ancient SARS-CoV epidemics may be important host determinants of SARS-CoV-2 (Souilmi et al., 2021).

Here, we aimed to identify key SARS-CoV adaptive loci and functional genomic differences between bats, which include the natural reservoir of SARS-CoVs, and primates, including humans. We performed high-throughput evolutionary and positive selection screens of 334 SARS-CoV-2 interacting proteins (Gordon et al., 2020) using the Detection of Genetic INNovation (DGINN) pipeline (Picard et al., 2020), followed by comprehensive functional-genetic analyses of seven VIPs of interest. We provide the results in the searchable VirHostNet 2.0 web portal. Using this approach, we identified 38 bat and 81 primate genes with strong evidence of positive selection. Of these, we found 17 proteins, including the ACE2 receptor, subjected to adaptative evolution in both clades, (i) confirming that past SARS-CoV epidemics occurred during both bat and primate evolution and (ii) identifying the core VIPs that shaped universal SARS-CoV*-*host molecular arms-races. We also identified 84 VIPs with lineage-specific adaptations that likely contributed to SARS-CoV pathogenicity in different mammalian hosts.

Among these, we uncover the important role of several genes, including TMPRSS2, FYCO1 or RIPK1 that play important roles in entry, trafficking or inflammatory responses, respectively. We hypothesize that these past adaptation events in bats and primates underlie differences in susceptibility to SARS-CoV-2 infections and key determinants in COVID-19 severity in modern humans.

## Results

### Characterization of the evolutionary history of SARS-CoV-2 VIPs in bats and primates

Because pathogenic viruses and hosts are engaged in evolutionary arms-races, adaptive signatures accumulate in VIP genes as a result of past epidemics (Daugherty and Malik, 2012; Enard et al., 2016). Adaptive evolution can be identified by positive selection analyses over a set of protein coding orthologs when their rate of non-synonymous codon substitutions exceeds that of synonymous ones (Sironi et al., 2015). To identify the proteins with such signatures of adaptive evolution, we studied the evolutionary history of the SARS-CoV-2 interactome identified in *in vitro* experiments. Furthermore, to discover key SARS-CoV-2 – host determinants of replication and pathogenesis, we aimed to identify the common and different evolutions and genetics of the VIPs in the human versus the reservoir host. We therefore performed comparative phylogenetics of the VIPs in primates and bats. Specifically, we studied the 332 host proteins identified by Gordon et al. in mass-spectrometry assays of SARS-CoV-2 proteins in human cells (Gordon et al., 2020), in addition to the angiotensin converting enzyme 2 (ACE2) receptor and the transmembrane protease serine 2 (TMPRSS2), both necessary for virus entry into the cells.

To perform the phylogenetic and positive selection analyses, we used the Detection of genetic innovation DGINN bioinformatic pipeline (Picard et al., 2020) that entirely automates the analyses and combines several methods to test for selection across large datasets (Figure 1A). Briefly, from each of the 334 human reference gene sequences, DGINN automatically retrieved bat and primate homologs (from NCBI nr database), curated the coding sequences, and performed a codon-alignment followed by a gene phylogenetic reconstruction (Figure 1A, Table S1). The pipeline then screened for duplication events and identified orthologs and potential paralogs, as well as recombination events. This mainly allows correct phylogenetic and positive selection analyses of VIPs from gene families, and with recombination events. Finally, each aligned set of orthologs was used to measure rates of codon substitutions and to estimate whether the whole gene, as well as any codon, are evolving under positive selection. For this, DGINN uses a combination of methods from the following selection tools: HYPHY (BUSTED and MEME), PAML (Codeml M0, M1, M2, M7, M8, and associated Bayesian Empirical Bayes (BEB) for codon-specific analyses), and bpp (M0^NS^, M1^NS^, M2^NS^, M7^NS^, M8^NS^, and associated Posterior Probabilities (PP) for codon-specific analyses) (Figure 1A, Methods for details, Picard et al 2020) (Guéguen et al., 2013; Pond et al., 2005; Yang, 2007).

**Figure 1.**
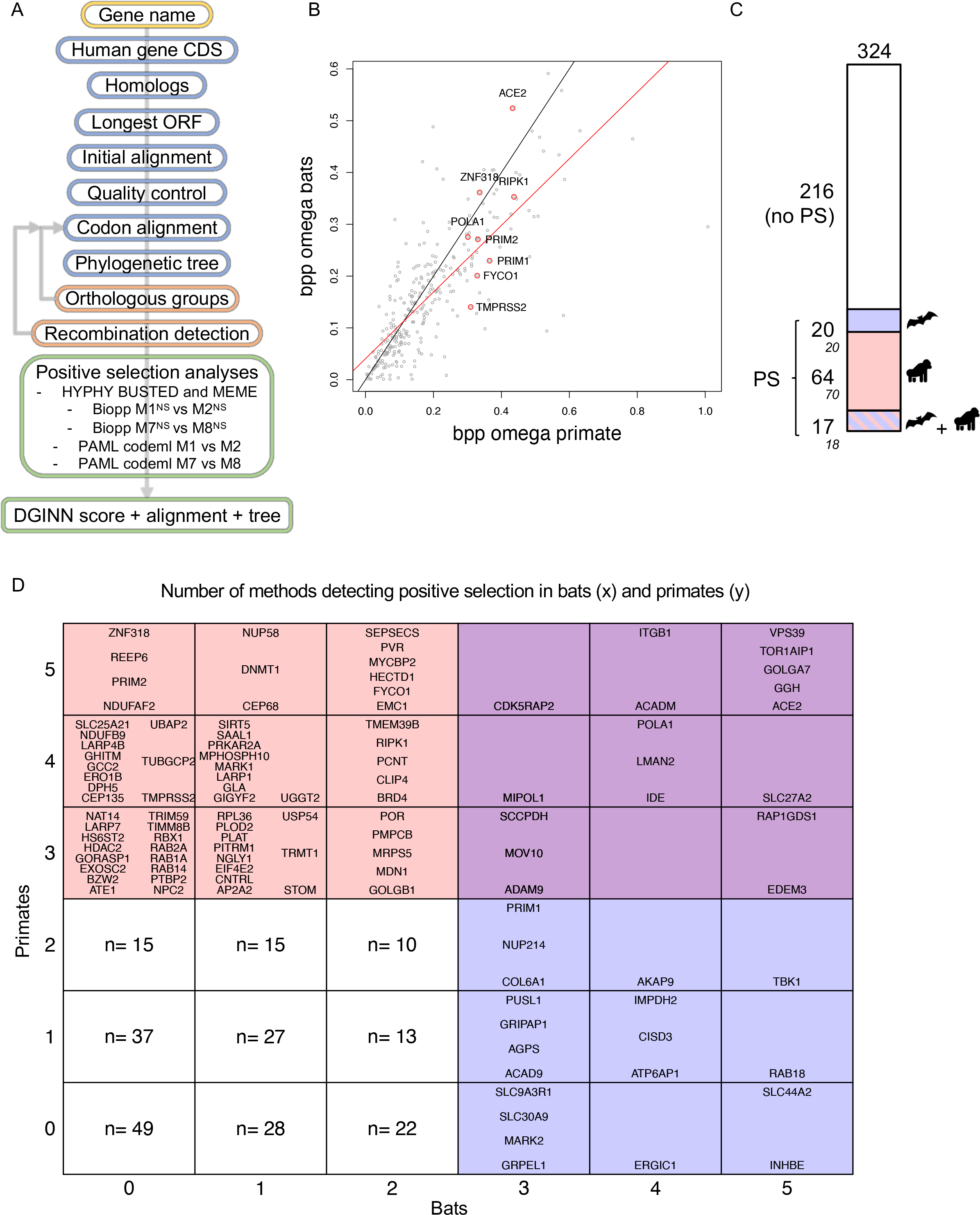
Identification of the SARS-CoV-2 interactome with signatures of positive selection in bats and primates. A, Overview of the DGINN pipeline to detect adaptive evolution in SARS- CoV-2 VIPs. B, Natural selection acting on bat and primate VIP genes. Comparison of omega (dN/dS) values of the VIPs during bat (y axis) and primate (x axis) evolution, estimated by bpp Model M0. In black, the bisector. In red, the linear regression. The names correspond to genes that we comprehensively analyzed (Table 1). C, Overview of the number of VIPs under significant positive selection (i.e., by at least three methods in the DGINN screen) in bats and/or primates. A total of 324 genes could be fully analyzed in the two mammalian orders. Numbers represent the number of genes in the categories: No PS (positive selection) or PS, within each host – represented by a pictogram. The numbers correspond to the conservative values after visual inspection of the positively selected VIP alignments, while the italic numbers are from the automated screen. D, Table showing the genes identified by x,y DGINN methods in bats and primates, respectively. For the genes with low DGINN scores (<3), only the number of genes in each category is shown (see Figure S4 for details). Of note, seven primate genes are “false positive”: EMC1 (ER membrane protein complex subunit 1), MOV10 (Mov10 RISC complex RNA helicase), POR (cytochrome p450 oxidoreductase), PITRM1 (pitrilysin metallopeptidase 1), RAB14, RAB2A, and TIMM8B (translocase of inner mitochondrial membrane 8 homolog B).

**Table 1.**
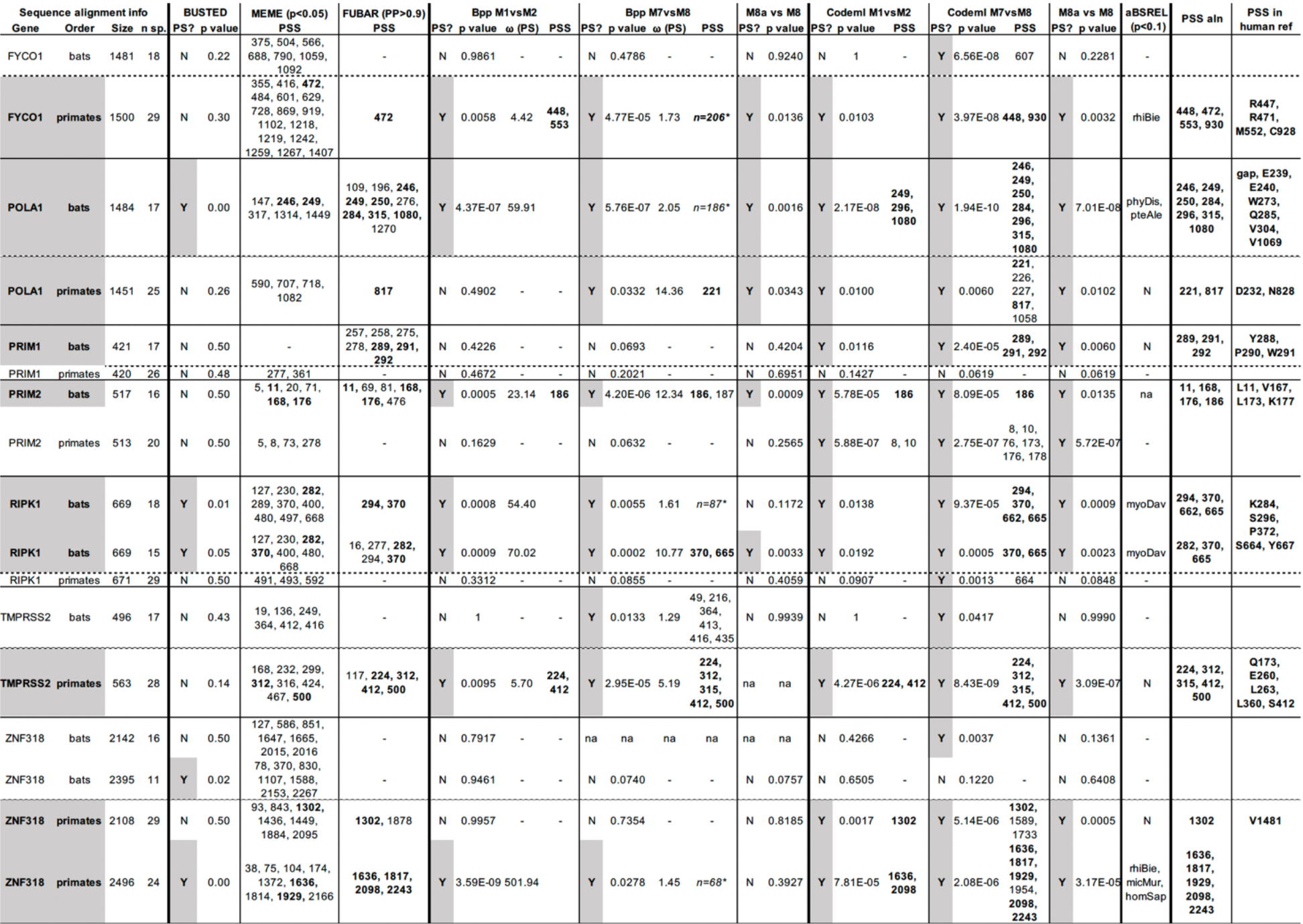
Results from the comprehensive positive selection analyses of the genes of interest. For each gene, are presented the results of the comprehensive phylogenetic and positive selection analyses: BUSTED, MEME, FUBAR, aBSREL from HYPHY/Datamonkey.com, M1vsM2, M7vsM8, M8avsM8 from Bpp, and M1vsM2, M7vsM8, M8avsM8 from PAML Codeml. The genes identified under positive selection are highlighted in grey. The sites considered under positive selection after the analyses are in “PSS aln” and “PSS in human ref”, corresponding to the site number in the codon alignment and the corresponding amino acid site in the human reference sequence. Alignments, trees, and interactive table are available at: https://virhostnet.prabi.fr/virhostevol/. Legend details: Size, length of the codon alignment; n. sp., number of species included in the alignment; PS?, if the gene is under positive selection: Y, yes, N, no; p value, supporting a model under positive selection; PSS, positive selection sites; the cutoff for each method is given in the table; omega (PS), corresponds to the omega value in the positive selection class (dN/dS>1). ZNF318 and the proteins from the Primase complex are in Supplementary Information, and in Figures S9 and S10, respectively. *, for bpp M8 PSS analyses there were dozens of sites under positive selection due to the low omega value in the class w >1. For aBSREL, the branch identified under positive selection is given by the DGINN nomenclature (three letters from the genus and three from the species). na, not available.

We found that the DGINN pipeline, previously validated on nineteen primate genes (Picard et al., 2020), was efficient at screening hundreds of genes and at analyzing other mammalian orders (here, chiroptera) (Figures S1-S2). Overall, our bioinformatic screen allowed us to obtain the bat and primate evolutionary history of 324 common SARS-CoV-2 VIP genes (i.e. 330 in bats and 329 in primates). We compiled the resulting sequence alignments, phylogenetic trees, and gene and site-specific positive selection results to an open-access and searchable web application (https://virhostnet.prabi.fr/virhostevol/), which constitutes a new public resource to visualize and download the evolution of SARS-CoV-2 VIPs in primates and bats.

### Identification and comparative analysis of SARS-CoV-2 VIPs with signatures of positive selection during bat and primate evolution

To characterize the overall trend in the evolution of each VIP in primates and in bats, we compared their omega parameter, which is positively correlated with the natural selection acting on the protein (Figure 1B). We found a similar trend in the natural selection of bat and primate genes: those with an elevated omega in primates had an overall rapid protein evolution in bats too. Beyond this trend, we cannot compare the omega values *quantitatively* between the two mammalian orders – reasons include differences in the number of analyzed species (i.e. 12 and 24 median number of species in bats and primates, respectively), the population sizes, the genetic distances, etc.

We next identified the genes with evidence of positive selection by at least three methods in the DGINN screen. In bats, we found 38 genes, roughly 12% of SARS-CoV-2 interacting proteins, with signatures of positive selection (Figure 1C). These include the ACE2 receptor, also reported by others as under strong positive selection in bats (Demogines et al., 2012; Frank et al., 2020). In primates, we identified 81 genes under positive selection, after discarding seven due to low-quality alignments and inclusion of erroneous sequences in the automatic steps (Figure 1D legend).

In the case of primate analyses, we identified more VIPs under positive selection than Gordon *et al*., in which they identified 40/332 genes under positive selection in primates using Codeml M8 vs M8a model (Gordon et al., 2020). One example is the Zinc finger protein ZNF318 that has some marks of positive selection during primate evolution in our analyses (Supplementary Information). However, the overall dN/dS estimate for each gene was highly similar between the two studies (Figure S3A) and we detected most of the genes they identified under positive selection: 38/40 VIPs under positive selection in Gordon *et al* also detected by ⩾1 DGINN method, including 28 by ⩾3 DGINN methods (Figure S3B-C). Thus, the main advantages of DGINN were the end-to-end automatic pipeline and the combination of multiple methods, thereby increasing sensitivity and specificity of positive selection analyses in screening approaches.

Altogether, we found 81 primate VIPs and 38 bat VIPs with evidence of positive selection (Figure 1C-D). Beyond Gordon *et al*., other SARS-CoV-2 *in vitro* and clinical studies also identified many of these positively selected genes as SARS-CoV-2 VIPs, thus confirming their suspected role as SARS-CoV-2 regulators or interacting proteins (Figure S5, (Parkinson et al., 2020)). Analyses of pathway enrichment showed that positively selected VIPs are strongly associated with cell cycle control and centrosome behavior biological pathways (Figure S6), suggesting that the control of cell division, and perhaps centrosome-regulated cell polarization, are important for SARS coronavirus *in vivo*.

We found 17 rapidly evolving genes shared between bats and primates, corresponding to 16% of all SARS-CoV-2 VIPs with evidence of positive selection (i.e., 17 genes in common over 108 in total) (Figure 1C-D). This list notably includes the ACE2 receptor of SARS coronaviruses that has undergone positive selection in both primates and bats (Figure 1B,D). It also includes known drug targets, such as the metalloprotease ADAM9 (Carapito et al.), the ITGB1 integrin (Sigrist et al., 2020), and POLA1 from the Prim-Pol primase complex (Chaudhuri, 2021) (Supplementary Information) (Figure 1D). Therefore, these genes may represent the core SARS-CoV VIPs that have been subjected to positive selection pressure during both primate and bat evolution.

However, we also identified 84 genes that have evolved through distinct selective pressures during primate and bat evolution – being under positive selection only in primates (64 VIPs) or bats (20 VIPs) (Figure 1C-D) – including TMPRSS2, FYCO1, RIPK1, ZNF318 and the Prim-Pol primase complex (Supplementary Information) that we will focus on. These genes represent VIPs with different evolutionary trajectories in bats and primates.

### Several SARS-CoV-2 VIPs under positive selection are VIPs of other coronaviruses and may also be interconnected with other viral families

To investigate whether the SARS-CoV-2 VIPs under positive selection are also known to interact with other coronaviruses, we interrogated the VirHostNet database (Guirimand et al., 2015) for interconnection with SARS-1 and MERS (beta coronaviruses), and CoV-NL63 and CoV-229 (alpha coronaviruses). We found 58 genes (i.e. 54% of 108 genes under positive selection in bats or primates) that are adaptive SARS-CoV-2 VIPs and also known interacting proteins of at least another coronavirus (Figure 2A). The positive selection marks in these VIPs therefore likely represent adaptations on host proteins that have regulated or interacted with coronaviruses over million years of coevolution with mammals. These coronavirus VIPs therefore represent an evolutionarily common set of coronavirus interacting proteins.

**Figure 2.**
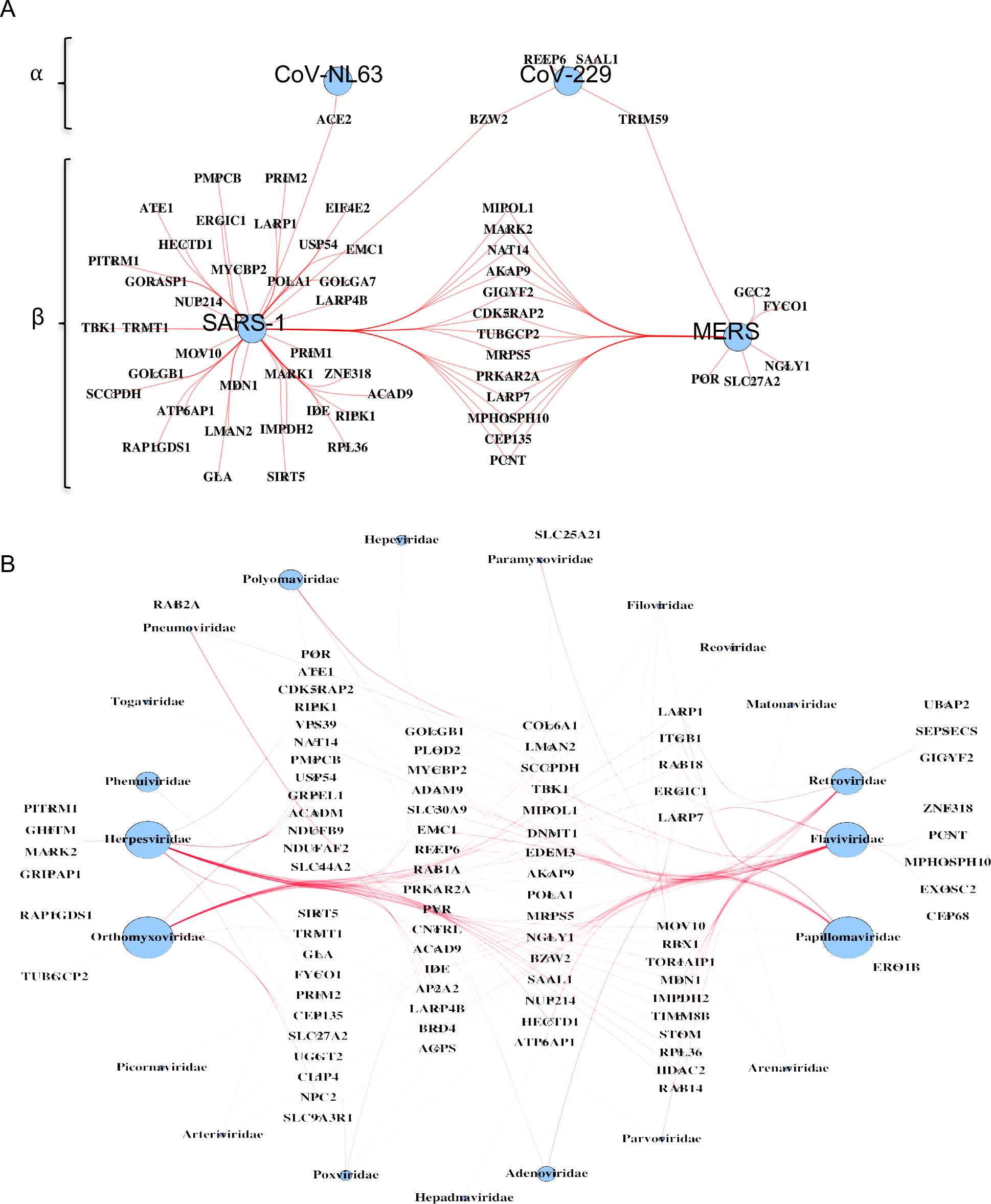
SARS-CoV-2 VIPs under positive selection are interacting proteins of other coronaviruses, as well as other viral families. Virus-host protein-protein interactions’ network of VIP genes under positive selection and interconnected with (A) other coronaviruses (from alpha- or beta-coronavirus genus), and (B) viral families other than coronaviruses. VIPs interacting with more than one additional viral family are in the center and arranged in columns (from left to right, interconnected with 2-6 different viral families). Node sizes at the virus families are proportional to the number of edges. The VIPs not interconnected are shown in Table S1.

Because positive selection may be driven by several viruses (Mitchell et al., 2013), we similarly investigated whether rapidly evolving SARS-CoV-2 VIPs were also functionally linked to other viral families (Figure 2B). We found that 82% of them (89 of 108 genes under positive selection in bats or primates) interconnected with one or more additional viral families beside coronaviruses. A number of proteins, including LARP1 and LARP7, ITGB1, Rab18 and ERGIC1, interconnected with six distinct viral families, highlighting their likely involvement as broad co- factors of viral replication (Figure 2B). On the other hand, several genes, such as FYCO1, ZNF318 or TMPRSS2, are interconnected with only 1-2 other viral families and may therefore represent more specialized VIPs (Figure 2B). Of note, although the TMPRSS2 co-entry factor has no other interactor in this analysis (Figure 2B, Table S2), it is a host factor for influenza virus entry (Böttcher et al., 2006; Limburg et al.). Lastly, the ACE2 receptor and other genes (Table S2) were not known to interact with other viruses, and therefore likely represent coronavirus-specific VIPs (Table S2).

### The SARS-CoV-2 predicted interface in TMPRSS2 has evolved under adaptive evolution in primates, but not in bats

Although the intrinsic role of TMPRSS2 in the cell is poorly known, this serine protease is a key factor for the cellular entry of SARS-CoV-2. TMPRSS2 is responsible for the priming of the viral spike S protein, an essential step for the ACE2 receptor recognition and the plasma cell membrane fusion process (Figure 3A) (Bestle et al., 2020; Hoffmann et al., 2020). In addition to SARS-CoV-2, other coronaviruses, including HCoV-229E, MERS-CoV, SARS-CoV-1, enter human cells in a TMPRSS2-dependent manner (Bertram et al., 2013; Iwata-Yoshikawa et al., 2019; Matsuyama et al., 2010). Whilst the genetic and functional adaptation of ACE2 has been studied (Demogines et al., 2012; Frank et al., 2020), the genetic diversification of mammalian TMPRSS2 is currently unknown. Our screen identified positive selection in TMPRSS2 in primates, but not bats, indicating that its functional diversification is specific to coronavirus adaptation in primates.

**Figure 3.**
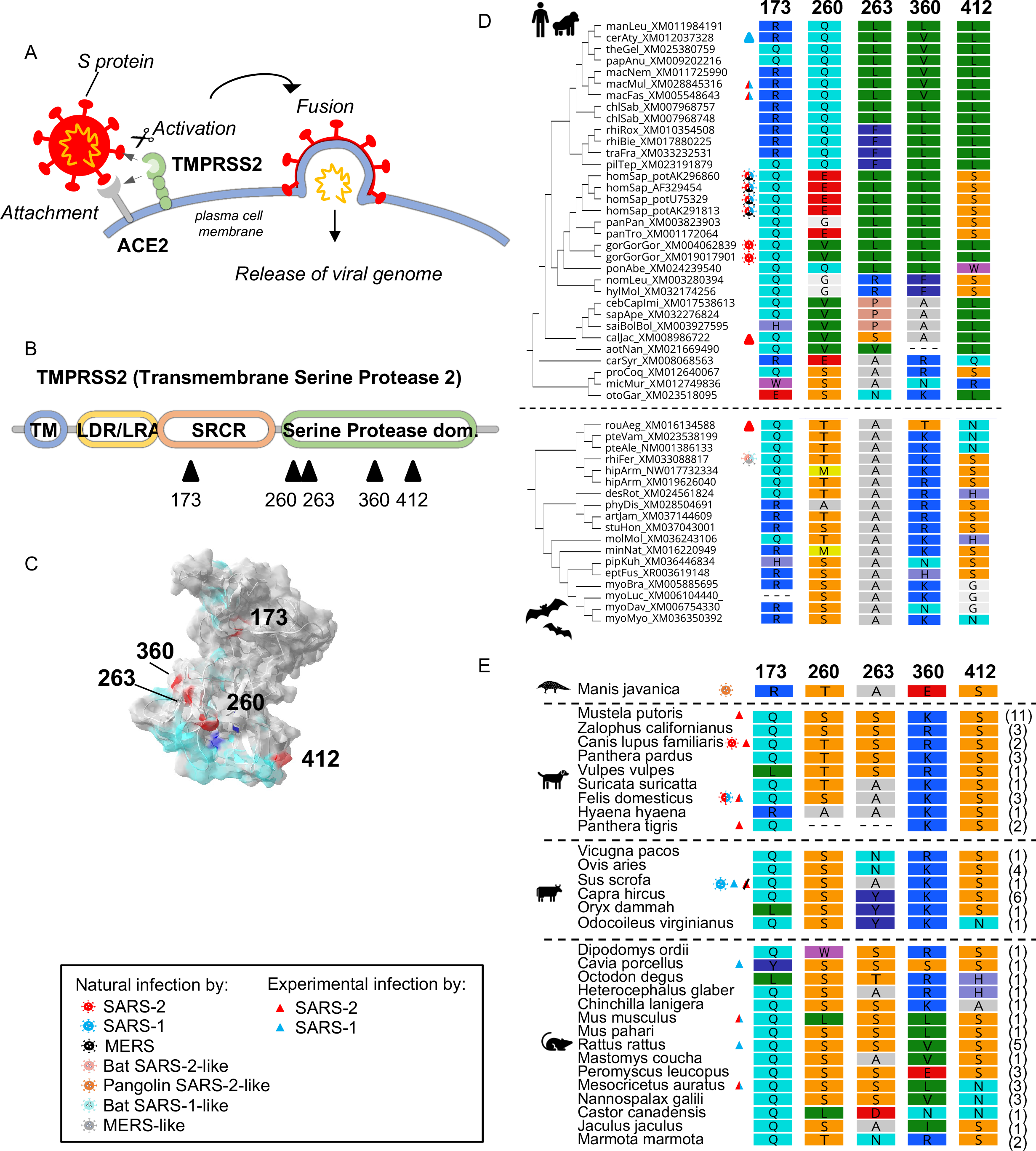
TMPRSS2 has evolved under strong positive selection in primates, but not in bats. A, Role of TMPRSS2 in SARS-CoV-2 entry. B, Diagram of TMPRSS2 predicted domains, with sites under positive selection in primates represented by triangles (Table 1). Codon numbering based on *Homo sapiens* TMPRSS2. C, 3D-structure modeling of human TMPRSS2 (amino acids 1-492) with the positively selected sites (red), the SARS-CoV-2 predicted interface (light blue), the catalytic site (dark blue). D, The positively selected sites identified in primate TMPRSS2 are highly variable in primates (top), but more conserved in bats (bottom) where they are not identified as under adaptive evolution. Left, cladograms of primate and bat TMPRSS2 with species abbreviation and accession number of sequences. Amino-acid color-coding, RasMol properties (Geneious, Biomatters). Icon legend is embedded in the figure, with multicolored pictograms/triangles showing cases fulfilling multiple conditions. E, Positively selected sites in primates exhibits different patterns of variability in other mammals: pangolin, carnivores, artiodactyls and rodents. Right, numbers in brackets correspond to the number of species within the order with the same TMPRSS2 haplotype at these positions (e.g. the QSSKS motif in *Mustela putoris* was found in eleven rodent species). The corresponding motif in species/cells susceptible or permissive to coronaviruses is shown in Figure S7.

To validate the screen results and further characterize TMPRSS2 evolution in both orders, we obtained sequences from additional primate and bat species that were not included in the automated DGINN screen. We therefore obtained two new high-quality codon alignments of TMPRSS2: from 18 bat species and from 33 primate species (https://virhostnet.prabi.fr/virhostevol/ “Genes of focus”, Table 1). From these comprehensive alignments, we first confirmed that TMPRSS2 has experienced significant and strong positive selection during primate evolution (Bio++ and PAML codeml M1 vs M2 p-values: 0.0095 and < 4.27 10^-06^, respectively). This was in contrast to its evolution in bats, in which we did not find evidence of selective pressure (Bio++ and codeml M1 vs M2 p-values: 1 and 1, respectively) (Table 1).

To identify the precise residues that have diversified during primate evolution, we performed site-specific positive selection analyses. We identified five residues (173, 260, 263, 360 and 412 – numbering from the human TMPRSS2 sequence) that were significantly detected under positive selection by at least two independent methods (Table 1, Figure 3B). Of note, position 197, which is polymorphic in human TMPRSS2 (rs12329760, V197M) and may be associated with COVID-19 severity ((Jeon et al., 2021) p-value around 10^-5^ above the 10^-8^ significance threshold commonly used in GWAS multiple testing), encoded for a conserved valine in all non-human primate sequences. Because the SARS-CoV-2 – TMPRSS2 interface is currently unknown, only *in silico* molecular docking studies have predicted the substrate binding region (Brooke and Prischi, 2020; Hoffmann et al., 2020; Rangel et al., 2020; SENAPATI et al., 2021). Remarkably, the sites under positive selection cluster nearby or within the predicted SARS-CoV-2-host interface (Figure 3C), suggesting that SARS-CoVs played a significant role in TMPRSS2 diversification. These regions of TMPRSS2 are also the target of several drugs, such as α1-antitrypsin (α1AT), Camostat mesylate, Nafamostat and Bromhexine hydrochloride inhibitors (Hoffmann et al., 2020; Li et al., 2021; Wettstein et al., 2021) and newly reported N- 0385 (Shapira et al., 2022), and could therefore be prioritized in functional studies.

Finally, by analyzing the physicochemical nature of the positively selected sites, we found that they encode for residues with very different properties, which would significantly impact the TMPRSS2 protein structure over primate evolution and lead to species-specificity at the virus-host interface. In particular, variation at key residues 260 and 412 was particularly high in Hominoids, but low in Old World monkeys (Figure 3D), suggesting lineage-specific adaptations within primates. To determine whether this domain of TMPRSS2 has been rapidly evolving in other mammals, we extended our analyses by retrieving other mammalian sequences. We found that most of these sites were overall conserved, except in rodents which exhibited high variability at positions 263 and 360 (Figure 3E). In bats, although none of the models identified significant positive selection in TMPRSS2, the sites 260, 360 and 412 were also variable (Figure 3D).

Comparing the variability between resistant and susceptible (naturally or experimentally) species to SARS-CoVs and MERS-CoVs did not reveal any clear pattern (Figure 3E). However, the location and extreme variability of the positively selected sites appear lineage-specific across mammals (with high amino acid toggling in some clades and conservation in others) and suggest that these residues, combined with ACE2 receptor variability, may contribute to SARS-CoV susceptibility and species-specificity.

Altogether, our findings support that the positive selection signatures in TMPRSS2 are reminiscent of ancient SARS-CoV-driven selective pressures during primate evolution.

Mutagenesis studies of TMPRSS2, guided by the evolutionary analyses, are now required to identify the exact and relevant SARS-CoV determinants, as well as the functional implication of the interspecies variability in TMPRSS2.

### Evidence that FYCO1 is involved in SARS coronavirus pathogenesis or replication at different time scales during primate evolution

FYCO1 (FYVE and coiled-coil domain containing 1) is involved in microtubule transport and autophagy (Figure 4A). Autophagy is an important degradation process of cytoplasmic proteins and organelles, which may be dysregulated during aging, diseases, and by pathogens. FYCO1 acts as an adaptor protein allowing the microtubule transport of autophagosomes in a STK4- LC3B-FYCO1 axis (Cheng et al., 2016; Nieto-Torres et al., 2021). Mutations of the human *FYCO1* gene cause autosomal-recessive congenital cataract, a major cause of vision dysfunction and blindness (Chen et al., 2011; Satoh et al., 2021). Until the COVID-19 pandemic, there was no report of FYCO1 involvement in viral infection. However, FYCO1 is among the very few genes identified in human genome-wide association studies (GWAS) to be significantly associated with severe COVID-19 (Pairo-Castineira et al., 2021; The Severe Covid-19 GWAS Group, 2020)(The COVID-19 Host Genetics Initiative, 2020). GWAS correlates natural genetic variants in human populations to phenotypic traits; here COVID-19 severity. Therefore, genes identified in GWAS may directly be involved in SARS-CoV-2 replication or pathogenesis.

**Figure 4.**
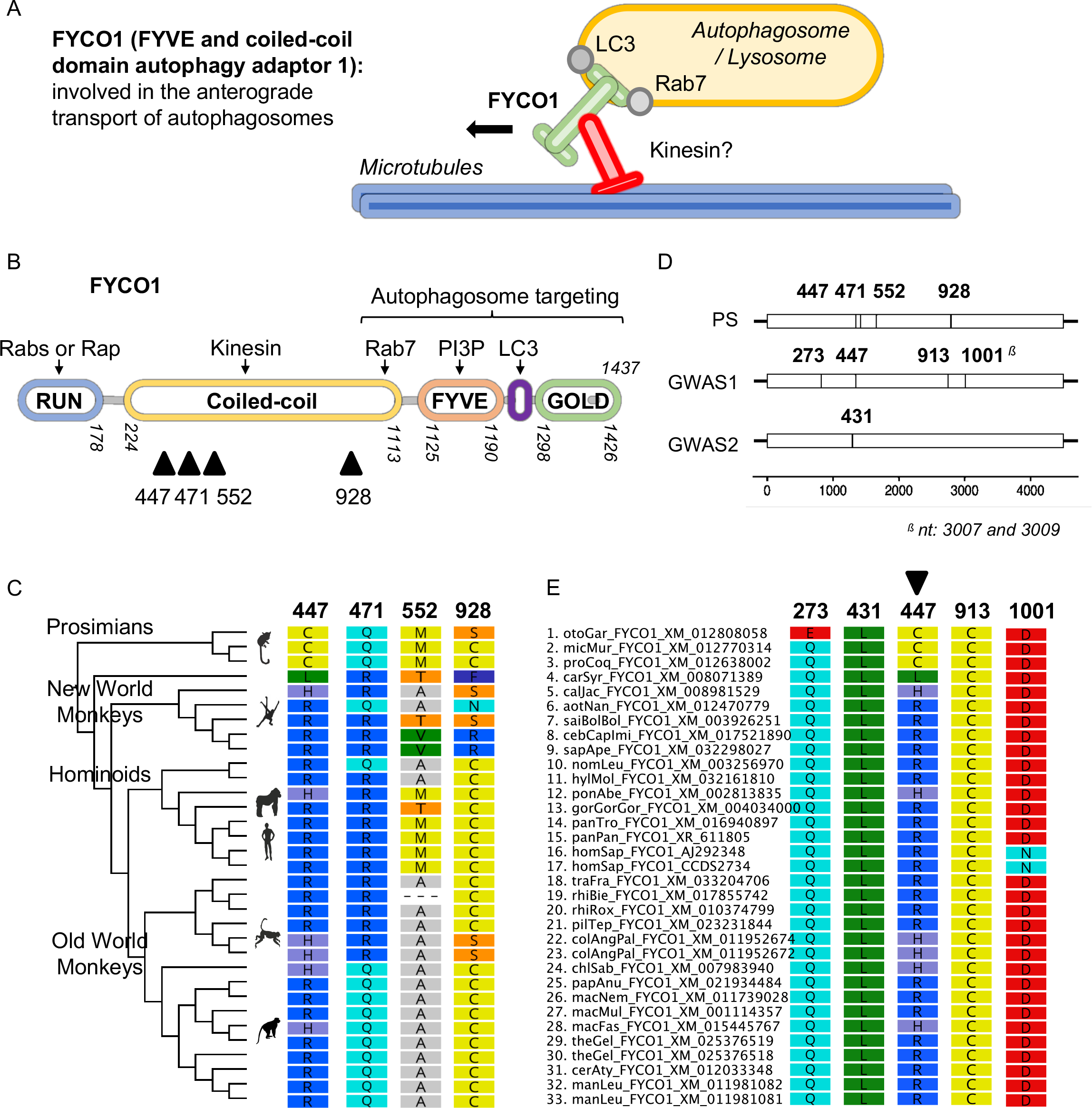
Domains of FYCO1 that are associated with severe COVID-19 in human have also evolved under significant positive selection in primates, but not in bats. A, Known cellular role of FYCO1. B, Diagram of FYCO1 predicted domains, with sites under positive selection in primates represented by triangles (Table 1). Codon numbering based on *Homo sapiens* FYCO1. C, Amino acid variation at the positively selected sites in primates. Left, cladogram of primate FYCO1 with major clades highlighted. The exact species and accession number of sequences are shown in Panel E. Amino-acid color-coding, RasMol properties (Geneious, Biomatters). D, Sites identified in the coding sequence of FYCO1 as under positive selection (PS) in primates (top) and as associated with severe COVID-19 in human from two GWAS studies (middle: GWAS1, COVID-19 Host Genetics Initiative, 2021; bottom: GWAS2, Pairo-Castineira et al. 2020). x axis, nucleotide numbering. E, Amino acid variations in primate species at the sites associated with severe COVID-19 in GWAS.

Furthermore, FYCO1 had a high MAIC score (Figure S5B, (Parkinson et al., 2020)), indicating that several studies suspect its involvement in SARS-CoV-2 pathogenesis or replication, including Gordon and colleagues that identified human FYCO1 interaction with SARS-CoV-2 NSP13 (Gordon et al., 2020).

As for TMPRSS2, the DGINN screen identified signatures of positive selection in primate FYCO1, but not in bat FYCO1. We then retrieved all FYCO1 sequences available for primates (29 species) and bats (18 species) and performed comprehensive phylogenetic and positive selection analyses. This new comprehensive positive selection analyses confirmed that FYCO1 has undergone positive selection in primates, but not in bats (Table 1).

Site-specific selection analyses identified four residues with strong evidence of significant positive selection in primates in at least two independent methods: 447, 471, 552, and 928 (Figure 4B-C, Table 1). Though no crystal structure is available for full-length FYCO1, these rapidly evolving sites fall into the coiled-coil domain of FYCO1, which is important for interaction with Kinesin. In addition, the different primate species encode for amino acids with very different physicochemical properties at these sites (Figure 4C), indicating potential structural and functional plasticity in this region. These positive selection marks may therefore represent virus-host interplays and be the result of selective pressure by ancient epidemics during primate evolution.

To correlate primate natural genetic variants with ongoing human polymorphisms and association with COVID-19 severity, we compared FYCO1 variations in primates with the human polymorphisms associated with increased SARS-CoV-2 pathogenicity (GWAS). Using the COVID-19 Host Genetics Initiative data (https://www.covid19hg.org/results/r6/) as well as the data from Pairo-Castineira and colleagues (https://genomicc.org/data/), we identified five codons in FYCO1 with polymorphisms associated with severe COVID-19 in humans (Figure 4D). By comparing these positions to the four positively selected sites in primates, we found one position in common, site 447 (genome position 45967996) (Figure 4C-E). This shows that residue 447, whose alleles are correlated with COVID-19 severity in human, has also been subjected to adaptive evolution in primate history. In addition, at the protein domain level, the regions 430- 555 and 910-1005 both have several residues associated with severe COVID-19 in humans and residues under adaptive evolution in primates (Figure 4D-E). Therefore, our combined positive selection and GWAS analysis identified FYCO1 regions that may be key host determinants of SARS-CoV-2 and COVID-19.

Overall, our results support the importance of FYCO1 in SARS coronavirus pathogenesis or replication in primates, in both ancient (our positive selection analysis) and modern (GWAS) times. Furthermore, observed differences in positive selection between the susceptible primate hosts and bats (where no positive selection was observed and no disease is known to be associated with CoV infection) may highlight key differences in pathogenesis. We have two main hypotheses for the role of FYCO1 in SARS-CoV infection. First, given its known cellular role (Figure 4A), FYCO1 may play a role in facilitating viral egress and replication. Second, FYCO1 may be involved in COVID-19 pathogenesis, potentially through an indirect mechanism by affecting the autophagy process or vesicle trafficking necessary to resolve viral infection.

### RIPK1 has been under adaptive evolution in bats at residues that are crucial for human RIPK1 regulation

Human RIPK1 is an adaptor protein involved in inflammation through the tumor necrosis factor alpha receptor 1 (TNFR1) and the Toll-like receptors 3 and 4 (TLR3/4), leading to pro-survival, apoptotic or necroptotic signals (Figure 5A) (Delanghe et al., 2020; Liu et al., 2018). A curated analysis of RIPK1 interactors showed that it is a central hub for 79 cellular partners involved in key inflammatory and cell survival/death processes (Reactome database; Figure S8A). RIPK1 interacts with SARS-CoV-2 NSP12 (RdRp) (Gordon et al., 2020), and is further involved in several bacterial and viral infections, being usurped by pathogens or involved in anti-microbial immunity (Figure S8B).

**Figure 5.**
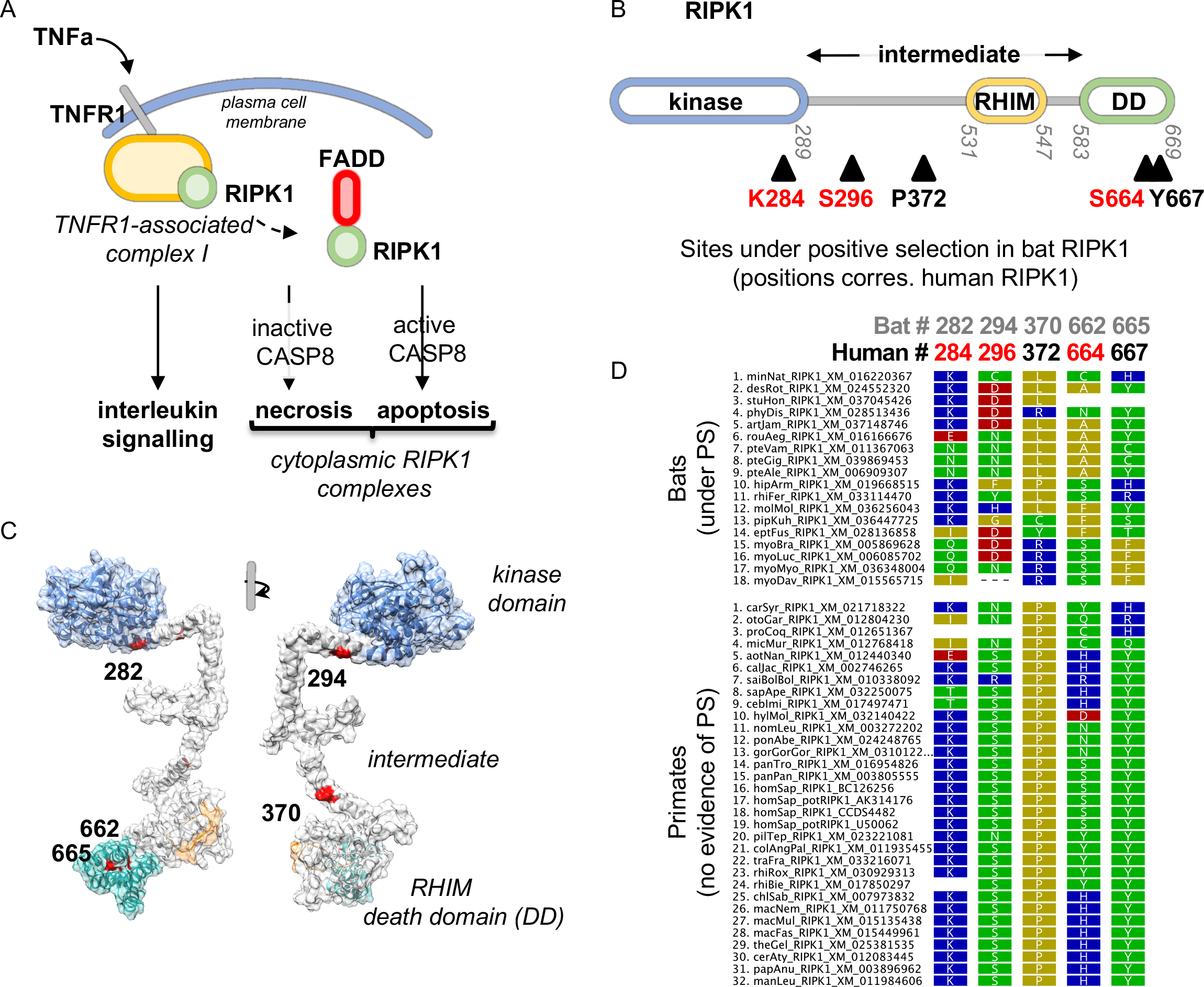
The multi-functional and inflammatory RIPK1 protein exhibits strong evidence of adaptation in bats at key regulatory residues. A, Schematic diagram of the three main functions associated to human RIPK1 in TNF signaling. As part of the TNFR1-associated complex, RIPK1 induces pro-survival signals that notably lead to NFkB activation. When dissociating from this complex, as a result of multiple events involving both phosphorylation and ubiquitination, RIPK1 can associate to FADD and lead to apoptosis or necrosis. B, Diagram of RIPK1 domains with the residues under positive selection in bats (black triangles) with the corresponding positions in human RIPK1 (Table 1). C, 3D-structure prediction of bat (*Rhinolophus ferrumequinum*) RIPK1, using RaptorX. The protein domains are color-coded as in B. Residues under positive selection are in red and numbered is according to their position in bat RIPK1. D, The positively selected sites identified in bat RIPK1 are highly variable in bats (top), but more conserved in primates (bottom), where they are not identified as under adaptive evolution. Left, bat and primate RIPK1 with species abbreviation and accession number of sequences. Amino-acid color-coding, Polarity properties (Geneious, Biomatters). The correspondence of residues from *Rhinolophus ferrumequinum* bat RIPK1 (grey) to human numbering (black) is shown at the top.

In our DGINN screens, we only identified signatures of positive selection in primate RIPK1. As previously, to obtain comprehensive phylogenetic and positive selection analyses, we retrieved all available coding sequences of bat (n=18 species) and primate (n=29 species) RIPK1 and performed new codon alignments and analyses. Here, we found strong evidence of positive selection in bat RIPK1, but not in primates (Table 1). This is different from our screen results, and this discrepancy was mostly due to (i) the addition of sequences as compared to our screens (i.e., from 12 to 18 bat species sequences, and from 24 to 29 primates) and (ii) the high-quality codon alignments, which are crucial for positive selection studies.

Next, using site-specific analyses, we identified five residues in bat RIPK1 that have evolved under significant positive selection (Figure 5B, Table 1). These are located in the intermediate domain (282, 294, 370) and in the C-terminal death domain DD (662, 665) of RIPK1. The latter domain can interact with other DD containing proteins, such as FADD, and has determinants for host-pathogen interactions (Delanghe et al., 2020; Liu et al., 2018). To determine where the positively selected sites fall in the three-dimensional protein, we used a structure prediction of bat RIPK1 from *Rhinolophus ferrumequinum,* which genus is a SARS-CoV-2 reservoir. We found that the rapidly evolving sites are exposed at the protein surface (Figure 5B-C; and Figure S8B for a comparison with the predicted 3D structure of human RIPK1, (Mompeán et al., 2018)). Therefore, physicochemical variations at sites 662 and 665 (Figure 5D) in the death domain could modulate interactions with DD-bearing proteins, and thus influence the ability of bat RIPK1 to drive cell death (Grimm et al., 1996). Alternatively, these variations may affect interactions between bat RIPK1 and viral antagonists, and thus may be directly involved in host-pathogen evolutionary conflicts.

Interestingly, using comparative analyses of bat and human RIPK1s, we found that the positively selected sites 282, 294 and 662 in bat RIPK1 correspond to sites K284 and S296, and S664 in human RIPK1, which are ubiquitinated and phosphorylated, respectively (Delanghe et al., 2020; Simpson et al., 2021) (Figure 5B in red). The posttranslational modifications at these sites are very important for the balance between the pro-survival and the pro-cell death functions of human RIPK1. It is thus possible that variation at these residues (Figure 5D) affects how bat RIPK1 is regulated.

Overall, our evolutionary analyses indicate that RIPK1 is an important SARS-CoV-2 (and other virus) interacting protein and suggest that residues undergoing positive selection in bats may be important (i) as determinants of virus-host interfaces, and (ii) as regulators of the protein balance between pro-survival and pro-cell death activities. The latter may allow certain bat species to tolerate viral infections and regulate the associated inflammation.

## Discussion

This study of the evolution of SARS-CoV-2 interacting proteins in mammals help us to understand how the bat reservoir and the primate host have adapted to past coronavirus epidemics and may shed light on modern genetic determinants of virus susceptibility and COVID-19 severity. Here, among the 334 genes encoding for SARS-CoV-2 VIPs, we identified 38 and 81 genes with strong signatures of adaptive evolution in bats and primates, respectively. Results are available at https://virhostnet.prabi.fr/virhostevol/. First, we found a core set of 17 genes, including the ACE2 receptor and POLA1, with strong evidence of selective pressure in both mammalian orders, suggesting (i) past epidemics of pathogenic coronaviruses in bats and primates shaping mammalian genomes, and (ii) common virus-host molecular and adaptive interfaces between these two mammalian host orders. This represents a list of host genes that should be prioritized and studied for roles in broad SARS-CoV replication. We also found several genes under positive selection only in bats or primates (such as RIPK1 or TMPRSS2), which highlight important differences in the coevolution of primate and bat with SARS-CoVs.

Furthermore, we discovered specific residues within the VIPs with typical marks of virus-host arms-races, which may point to precise SARS-CoV-host interfaces that have been important *in vivo* and may therefore represent key SARS-CoV-2 drug targets (such as TMPRSS2 or FYCO1). Finally, we found that FYCO1 sites with hallmarks of positive selection during primate evolution are those associated with severe COVID-19 in humans, supporting the importance these rapidly evolving residues in SARS-CoV-2 pathogenesis and replication. Overall, our study identified several host factors that (i) have been driven by ancient epidemics of pathogenic SARS coronaviruses, (ii) are different between the bat reservoir and the primate host, and (iii) may represent key *in vivo* virus-host determinants and drug targets.

The difference in adaptive VIPs in primates and bats suggests that beyond the common virus-host interfaces, SARS-CoVs have an intrinsically different interactome in these distant hosts (i.e. specialization). Therefore, SARS-CoVs may have adapted to usurp and/or antagonize different cellular proteins in the primates versus the bats. This is exemplified by the evolution of the entry factor TMPRSS2 (amongst others). We identified strong evidence of virus-host arms-races in primates, but not in bats, suggesting that SARS-CoVs may not strongly rely on TMPRSS2 for entry in bat cells, as opposed to primates. Only functional studies on SARS-CoV natural entry pathways into bat cells would firmly determine this. Interestingly, the recent SARS-CoV-2 Omicron variant has evolved to enter the human cell through a TMPRSS2-dependent and -independent route, showing also intra-host species plasticity at these interfaces (Meng et al., 2022; Peacock et al., 2022; Pia and Rowland-Jones, 2022; Willett et al., 2022). Lastly, the importance of lineage-specificity of SARS-CoV-2 VIPs has previously been highlighted for OAS1. Indeed, humans rely on prenylated OAS1 to inhibit SARS-CoV-2 replication and prevent COVID-19 severity (Soveg et al., 2021; Wickenhagen et al., 2021), but *Rhinolophidae* bats do not encode for an OAS1 capable to interact with SARS-CoV-2 (Wickenhagen et al., 2021). Thus, in addition to genes such as TMPRSS2, FYCO1, or RIPK1, our findings provide dozens of genes that represent host-specific interfaces and may be critical *in vivo* SARS-CoV VIPs.

The differences between primate and bat evolution of the SARS-CoV-2 interactome may further result from important differences in the adaptation at the virus-host interface in a reservoir host versus a recipient host. In this model, beyond the core SARS-CoV-2 interactome of bats and primates, the genes under positive selection would correspond to host-specific adaptations to SARS-CoV. This could underlie important immunomodulatory differences between primates and bats (Christie et al., 2021). For example, the inflammatory protein RIPK1 showed signatures of adaptive evolution in bat residues that correspond with loss of important RIPK1 regulatory phosphorylation and ubiquitination residues in humans. With the caveats that no functional studies exist on bat RIPK1, the extrapolation of the functions ascribed to the corresponding residues in human RIPK1 suggests that positive selection in bat RIPK1 may result from an advantageous decrease of RIPK1-driven inflammation in bats. This is analogous to the loss of S358 phosphorylation site in bat STING that participates in a dampened inflammation response in bats (Xie et al., 2018), and supports a model where hosts that are more tolerant to viral infection contribute to the establishment of a host reservoir, such as hypothesized for bats (Ahn et al., 2019; De La Cruz-Rivera et al., 2018; Irving; Pavlovich et al., 2018; Prescott et al.; Xie et al., 2018; Zhang et al., 2013).

It is also possible that there are fewer signatures of adaptation in SARS-CoV interacting proteins in bats over primates, because coronaviruses may have been less pathogenic in the former host, and therefore less selective (Emerman and Malik, 2010; Irving). However, evidence of strong positive selection in the bat ACE2 receptor driven by ancient pathogenic SARS-CoVs (this study, and (Demogines et al., 2012; Frank et al., 2020)) supports a model in which past SARS-CoV epidemics have been sufficiently potent to shape bat genomes.

Our work also tries to bridge studies of ancient and recent evolution of genes, which can help us better understand past epidemics and adaptive genes, and ultimately develop evolutionary medicine. This study over millions of years of evolution (at the inter-species level) shows evidence of very ancient epidemics of SARS-CoVs that have shaped both primate and bat genomes. Marks of adaptation in SARS-CoV-2 VIPs at the human population level further identified evidence of past SARS-CoV epidemics in more recent human history (Souilmi et al., 2021). Bridging these ancient and more recent evolutionary analyses with GWAS studies would bring more direct confirmation of the causal role of viral interacting proteins in pathogenesis.

This is here exemplified by the FYCO1 gene that may be a central protein in SARS-CoV-2 pathogenesis and disease etiology.

A limitation of our study is that we did not quantify the selective pressures occurring at (regulatory) noncoding regions of the VIPs. Using human population genomics, Souilmi et al found that marks of positive selection have been particularly strong at non-coding regions of SARS-CoV interacting proteins (Souilmi et al., 2021). However, these analyses are challenging at the interspecies level, and more methods and high-quality genome alignments would be necessary for state-of-the-art mammalian genomic analyses. Our findings are therefore conservative and other marks of adaptation in the same, and in more, VIPs are certainly at play.

At the heart of our study analyzing the coding sequences of SARS-CoV-2 VIPs is the identification of site-specific adaptations at multiple SARS-CoV-2 interacting proteins, which may reflect the exact sites of molecular arms-races of proviral and antiviral VIPs with SARS-CoVs (Duggal and Emerman, 2012; Sawyer et al., 2005; Sironi et al., 2015). These sites are therefore of primary importance to investigate in functional assays to firmly identify key SARS-CoV-2-cell determinants and drug targets. For example, our study highlights TMPRSS2 and RIPK1, amongst others, as potential targets of interest. Primidone, an FDA-approved RIPK1 inhibitor, has proven ineffective as a direct inhibitor of viral replication in established cell lines (Gordon et al., 2020; Riebeling et al., 2021). However, our findings suggest that RIPK1 inhibitors will more likely exert an effect on the virus-induced hyperinflammation, rather than on viral replication itself. As such, the evaluation of the effectiveness of RIPK1-kinase inhibitors will require a more complex cellular setup. Lastly, other viruses may also have driven adaptation at these VIPs, which therefore represent essential host-pathogens interfaces. Targeting the identified VIPs with strong marks of virus-host arms-races may be an effective broad antiviral strategy.

## Methods

### DGINN Screens

Analyses were performed as previously described in Picard et al., 2020. Briefly, CCDS identifiers were downloaded from HGNC Biomart (http://biomart.genenames.org/) for all 334 genes of interest. If there were more than one CCDS, the longest was selected. Initial codon alignments and phylogenetic trees were obtained using DGINN with default parameters (prank -F -codon; version 150803, HKY+G+I model (Löytynoja and Goldman, 2008); PhyML v3.2 (Guindon et al., 2010)). Duplication events were detected through the combined use of Long Branch Detection and Treerecs (Comte et al., 2020) as implemented in DGINN. Recombination events were detected through the use of GARD (Kosakovsky Pond et al., 2006) from the HyPhy suite as implemented in DGINN. For each VIP gene, the analyses of primate evolution and of bat evolution were separately run. The species trees employed for the tree reconciliation with Treerecs are accessible at: https://virhostnet.prabi.fr/virhostevol/. Positive selection analyses were then run using models from BUSTED and MEME from the HyPhy suite (Murrell et al., 2012, 2015; Pond et al., 2005) and codon substitution models from PAML codeml (M0, M1, M2, M7, M8) (Yang, 2007), and from Bio++ (M0, M1^NS^, M2^NS^, M7^NS^, M8^NS^) (Guéguen et al., 2013) as implemented in DGINN (Picard et al., 2020). For the chiroptera screen, a visual inspection of the resulting gene alignments was performed, and we refined 28 of them to delete erroneous ortholog sequences, erroneous isoforms, or sequencing errors. These 28 curated alignments were then re-analyzed with DGINN starting at the “alignment” step and included in the final results (Figure S1A, Table S1).

### MAIC Scores

MAIC scores were obtained from the database for COVID-19 (https://baillielab.net/maic/covid19/, 2020-11-25 release) (Parkinson et al., 2020). The 334 VIP genes were cross-referenced against the 10,000 best hits of the MAIC database.

### Detailed phylogenetic analyses on genes of interest

Alignments from the DGINN screens were retrieved and sequences that appeared erroneous were taken out. To obtain a maximum number of species along primate and bat phylogenies, further sequences were retrieved from NCBI databases using BLASTn. Final codon alignments were then made using PRANK (Löytynoja and Goldman, 2008) or Muscle Translate (Edgar, 2004), and phylogenetic trees were built using PhyML with HKY+I+G model and aLRT for branch support (Guindon et al., 2010).

Each curated gene alignment and tree were then submitted to positive selection analyses using the DGINN pipeline: HYPHY BUSTED and MEME, PAML Codeml (M0, M1, M2, M7, M8, M8a) and Bio++ (M0, M1^NS^, M2^NS^, M7^NS^, M8^NS^, M8a^NS^) (references in “DGINN screen”). To test for statistical significance of positive selection in Codeml and Bio++, we ran a chi-squared test of the LRT from models disallowing positive selection versus models allowing for positive selection (M1 versus M2, and M7 versus M8) to derive p-values. To identify the sites under positive selection, we used HYPHY MEME (p value < 0.05), the Bayes Empirical Bayes statistics (BEB) from the codeml M2 and M8 models (BEB > 0.95), and the Bayesian Posterior Probabilities (PP) from the M2^NS^ and M8^NS^ models in Bio++ (PP > 0.95). Three other web-based methods were used for this set of genes: A Fast, Unconstrained Bayesian AppRoximation for Inferring Selection (FUBAR) method to detect site-specific positive selection (PP > 0.90) (Murrell et al., 2013) and An adaptive branch-site REL test for episodic diversification (aBS-REL) to detect branch/lineage-specific positive selection (p-value < 0.1) (Smith et al., 2015).

### GWAS analyses

Using the COVID-19 Host Genetics Initiative data (https://www.covid19hg.org/results/r6/), we extracted the positions of human polymorphisms associated with “very severe respiratory confirmed covid vs. population” that are within FYCO1 coding sequence and have a p-value below 10^-8^. We similarly retrieved the positions found associated to severe COVID-19 by Pairo-Castineira et al. 2020 from the data publicly available at https://genomicc.org/data/ (Pairo-Castineira et al., 2021). We then matched the coordinates of polymorphic sites significantly associated with severe COVID-19 to the alignment of coding sequences of FYCO1 (using transcript FYCO1-205). To note, none of the other genes under positive selection contained polymorphism significantly associated with “very severe respiratory confirmed COVID” by the COVID-19 Host Genetics Initiative (see online browser).

### Reactome analyses

Gene pathway enrichment analyses were carried out on the Reactome biological pathways tools (https://reactome.org/). Interactors of RIPK1 were retrieved using the Reactome FIV plugin in Cytoscape (Gillespie et al., 2022).

### Protein structure predictions

Protein structure prediction of human and *Rhinolophus ferrumequinum* RIPK1 were modeled using RaptorX (Wang et al., 2017) and structures were visualized using the Chimera software (Pettersen et al., 2004).

Since no crystal structure is available for TMPRSS2 protease, the 3D structure of TMPRSS2 was predicted using the Iterative-Threading ASSEmbly Refinement (I-TASSER) server (Yang and Zhang, 2015). A total of 492 amino acid sequence of human TMPRSS2 obtained from NCBI Genbank (accession number AF329454) was used as query. The best model inferred by I- TASSER was selected using the C-Score – a measure assessing the quality of the models. The final estimates are: model C-score, -0.41; estimated TM-score, 0.66±0.13; RMSD, 8.2±4.5Å. The corresponding TMPRSS2 structure was generated using Swiss PDB viewer software (Johansson et al., 2012).

### Sequence logo generation

The amino acid sequence logos of TMPRSS2 were generated using WebLogo (V. 2.8.2, (Crooks et al., 2004)), based on an alignment of the positively sites from mammalian species reported as naturally susceptible and/or experimentally permissive to SARS-COV2, SARS-COV and MERS-COV.

### Code and Data availability

All codes are available in: https://gitbio.ens-lyon.fr/ciri/ps_sars-cov-2/2021_dginn_covid19, and the DGINN pipeline is available at: http://bioweb.me/DGINN-github. All results from the DGINN screens (“DGINN full dataset”) and from the detailed evolutionary analyses (“DGINN genes of interest”) are available through the VirHostEvol web service https://virhostnet.prabi.fr/virhostevol/. The Shiny web application open-source code is available from the IN2P3 gitlab https://gitlab.in2p3.fr/vincent.navratil/shinyapps-virhostevol.

## Acknowledgements

We sincerely thank Janet Young for sharing her raw data on the positive selection of primate genes for comparative analyses, as well as for her precious feedback on the project. We also thank Michael Emerman, Pierre-Olivier Vidalain, Dominique Guyot, and the members of LP2L for helpful discussions. We thank Sara Clohisey for generously providing us with the raw data for MAIC scores. We thank all the contributors of publicly-available genomic and genetic sequences, and of phylogenetic programs. Some molecular graphics and analyses were performed with UCSF Chimera, developed by the Resource for Biocomputing, Visualization, and Informatics at the University of California, San Francisco, with support from NIH P41-GM103311. This work was performed using the computing facilities of the PSMN (Pôle Scientifique de Modélisation Numérique) of the ENS de Lyon, the IFB (Institut Francais de Bioinformatique), and the CC LBBE/PRABI, and with the support from the BIBS (Bioinformatic and biostatistics service) of the CIRI, Lyon. We also thank S. Delmotte and B. Spataro for the cloud openstack.

Funding: LE and LG are supported by grants from the ANR LABEX ECOFECT (ANR-11-LABX- 0048 of the Université de Lyon, within the program Investissements d’Avenir [ANR-11-IDEX- 0007]) and the French Agence Nationale de la Recherche, under grant ANR-20-CE15-0020-01 (Project “BATantiVIR”). LE is further supported by the CNRS and by grants from the French Research Agency on HIV and Emerging Infectious Diseases ANRS/MIE (no. ECTZ19143 and ECTZ118944), and the CNRS France-U. Arizona Institute for Global Grand Challenges. AM is supported by the Fondation pour la Recherche Médicale, FRM:AJE201912009932.

## SUPPLEMENTARY

### Supplementary Information – Results and Discussion

#### ZNF318 has undergone positive selection at three sites in primates, but not in bats

ZNF318 (NP_055160, also known as TZF, testicular zinc finger protein) is a putative RNA binding Matrin-type zinc finger protein. Matrin-type zinc fingers are RNA interacting domains found most notably in matrins and U1 small nuclear ribonucleoprotein C (InterPro: IPR003604). Aside from two central matrin-type ZNF domains, ZNF318 interacts with the androgen receptor and forms homodimers through its N-terminal tail (Figure S9A) (Tao et al., 2006). Finally, despite accounting for about half the protein, the Proline-rich C-terminal domain of ZNF318 has currently no ascribed function (Figure S9A). While the precise molecular function of ZNF318 remains unknown, it was shown to interact with the HUSH chromatin silencing complex and contributes to splicing-coupled transposon, and potentially latent HIV, silencing (Chougui et al., 2018; Douse et al., 2020). Antiviral functions have also been described for some human matrin genes (Yedavalli and Jeang, 2011).

Our DGINN screens identified signatures of positive selection in ZNF318 from primates, but not in bat species. While we confirmed the presence of ZNF318 orthologs in 29 primate and 16 bat species, many were 5’ truncated. Thus, we used 5’ trimmed CDS for in-depth selection analyses (Table 1). Unlike the majority of codons that are highly conserved (mean dN/dS of ∼0.3), the C-terminal portion of ZNF318 displays many non-synonymous substitutions between orthologs. Statistical analyses using CodeML identified three sites under positive selection in this domain (M7 vs M8, BEB > 0.95): residues 1481, 1756 and 1908 in the corresponding human full-length protein (NP_055160, Figure S9, Table 1). There does not seem to be strong co- evolutionary signatures between these three residues and the combination of sites found in human (“V”, “R” and “P”) arose once in the last common ancestor with chimps and bonobos (Figure S9B). Of the three sites, non-synonymous substitutions altering the proline residue 1908 might have strongest structural impact. Interestingly, these residues show some level of variation in bats, with bias for Threonines and Alanines at orthologous positions of sites 1481 and 1908.

While the function of ZNF318 during SARS-CoV2 infection remains to be determined, our finding of positive selection in primates suggest that it contributed to host adaptation in this lineage. Moreover, our analysis identified ZNF318 Proline-Rich domain as a putative functional interface with viruses. Hence, it would be interesting to investigate if this domain carries functions relating only to viral infections, including with non-coronaviruses (Figure 2), or if it is also involved in ZNF318 other cellular functions.

#### Rapid evolution of the Prim-Pol primase complex (POLA1, PRIM1, PRIM2) in primates and bats

Initiation of DNA replication in eukaryotes is dependent on the multisubunit primase-polymerase alpha (Prim-Pol α) complex that is responsible for the *de novo* synthesis of RNA/DNA primers on both the leading and lagging strands. Primase consists of the small catalytic subunit PRIM1 (p49) and the large regulatory subunit PRIM2 (p58), while polymerase consists of the catalytic subunit POLA1 (p180) and the accessory subunit POLA2 (p70). During primer synthesis, the primase subunits generate short RNA oligos, which are then subsequently extended with DNA by Pol α to be further elongated by replicative DNA polymerases. While many viruses encode their own Prim-Pol (Guilliam et al., 2015), including a putative CoV primase consisting of NSP7 and NSP8, SARS-CoV-2 NSP1 interacted with all four subunits of Prim-Pol α (Gordon et al., 2020).

From our initial DGINN analysis, we identified positive selection on three out of the four Prim-Pol α subunits: POLA1 (four tests for primates and bats), PRIM1 (three tests for primates and two tests for bats), and PRIM2 (five tests for primates only). POLA2 was not identified under positive selection in primates or bats (one test for bats and zero tests for primates).

To validate the finding and to precisely characterize gene evolution in primates and bats for POLA1, PRIM1, and PRIM2, we retrieved additional sequences for these orders and generated new high-quality codon alignments. POLA1 still showed significant signatures of positive selection across both primate and bat lineages (Table 1). PRIM1 maintained signatures of positive selection in bats (Table 1). PRIM2 maintained signatures of positive selection in primates for some tests, and the additional sequences identified positive selection in bats not seen with our initial pipeline (Table 1). Together this confirmed our initial analysis that POLA1, PRIM1, and PRIM2 are rapidly evolving in primate and bat lineages, with bats having more robust positive selection than primates.

Our codon-specific positive selection analysis identified multiple residues that have rapidly evolved in primates and bats. We specifically focused on sites that were found with more than one test of selection (Table 1). To determine if these sites of positive selection were found at the interface between subunits of the Prim-Pol α complex or at surface exposed sites, we mapped PS sites onto the human Prim-Pol α crystal structure (PDB: 5EXR) (Baranovskiy et al., 2016). None of the sites were found at Prim-Pol α complex interfaces, and while many were clustered close together, all were surface exposed or found in putative unstructured regions that were not present in the crystal structures (Figure S10). Together, this indicates that the positive selection identified for this complex is not being driven by complex formation and intra-complex coevolution, or DNA replication.

POLA1 is of particular interest as it has been modeled to dock with SARS-CoV-2 NSP1 (PDB 7OPL)(Kilkenny et al., 2022) (Chaudhuri, 2021). However, none of the sites identified in our positive selection analysis were at the predicted POLA1-NSP1 interface (residues 615-629; Figure S10A-B), suggesting that NSP1 is not driving positive selection on POLA1. Instead, 7/9 PS sites in bats and primates were found in unstructured regions that were not present in the crystal structure, with four sites (V235, D232, E239, and E240) all falling in a predicted disordered region of the protein (amino acids 232-251). That this unstructured region showed strong positive selection in both primates and bats could suggest that primate and bat POLA1 rapid evolution is being driven by similar unknown pressures in these distinct families.

Altogether, our positive selection analysis has identified that the Prim-Pol α complex is under strong positive selection in bats and primates, however this is not driven by Prim-Pol α complex formation or NSP1. Thus, it remains unclear what is driving PS on the Prim-Pol α complex, whether SARS-CoV interact with the Prim-Pol α complex as a whole or individual proteins from this complex, and why SARS-CoV-2 directly interact with nuclear host DNA replication machinery.

So why does SARS-CoV-2 recruit the Prim-Pol α complex? One possibility is that Prim- Pol α may have a role in the innate immune response to viral infection, and thus viruses directly antagonize components of this complex. Prim-Pol α was identified to interact specifically with NSP1 (Gordon et al., 2020), which functions to inhibit host translation and innate immunity (Narayanan et al., 2008; Schubert et al., 2020). Pola1 is found in both the nucleus and cytoplasm, where it generates RNA-DNA hybrids that may be important for innate immune sensing (Starokadomskyy et al., 2016). Loss of function mutations in POLA1 lead to increased pathogen infection, innate immune activation, and decreased number and effectiveness of NK cells (Starokadomskyy et al., 2016)(Starokadomskyy et al., 2019). While this supports a role of POLA1 in innate immunity against pathogens, roles for PRIM1 and PRIM2 have yet to be investigated.

It is also possible that SARS-CoV are usurping the host primase complex (or components of this complex) to enhance genome replication. SARS-CoV NSP7 and NSP8 are proposed to function as primase important for initiation of genome replication (Konkolova et al., 2020; Zhai et al., 2005). However, recent cryoEM structures of NSP7-8-12 complex suggest that NSP7 and NSP8 are too far from the RdRP NSP12 active site to act as primase (Kirchdoerfer and Ward, 2019). Thus, it is possible that the host Prim-Pol α is recruited by NSP1 to further help initiation (and/or elongation) of CoV genome replication.

## Supplementary Figures

**Figure S1.**
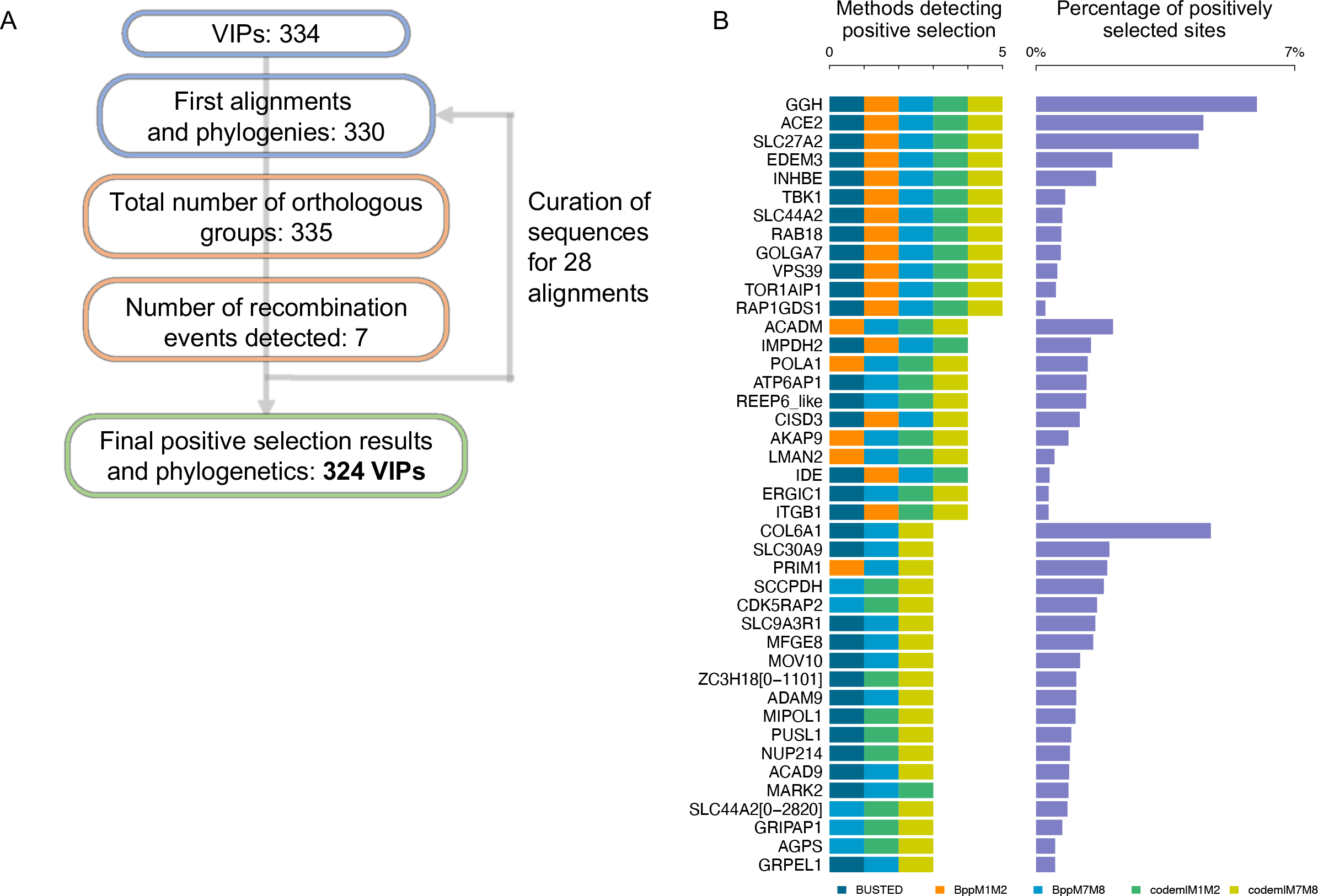
Identification of the SARS-CoV-2 interactome with signatures of positive selection in bats. A, Overview of the key steps of the bat VIP DGINN screen workflow. Details in Table S1. B, VIP-encoding genes identified under significant positive selection by at least three methods in DGINN (embedded legend). The percentage of positively selected sites in each VIP is shown on the right panel.

**Figure S2.**
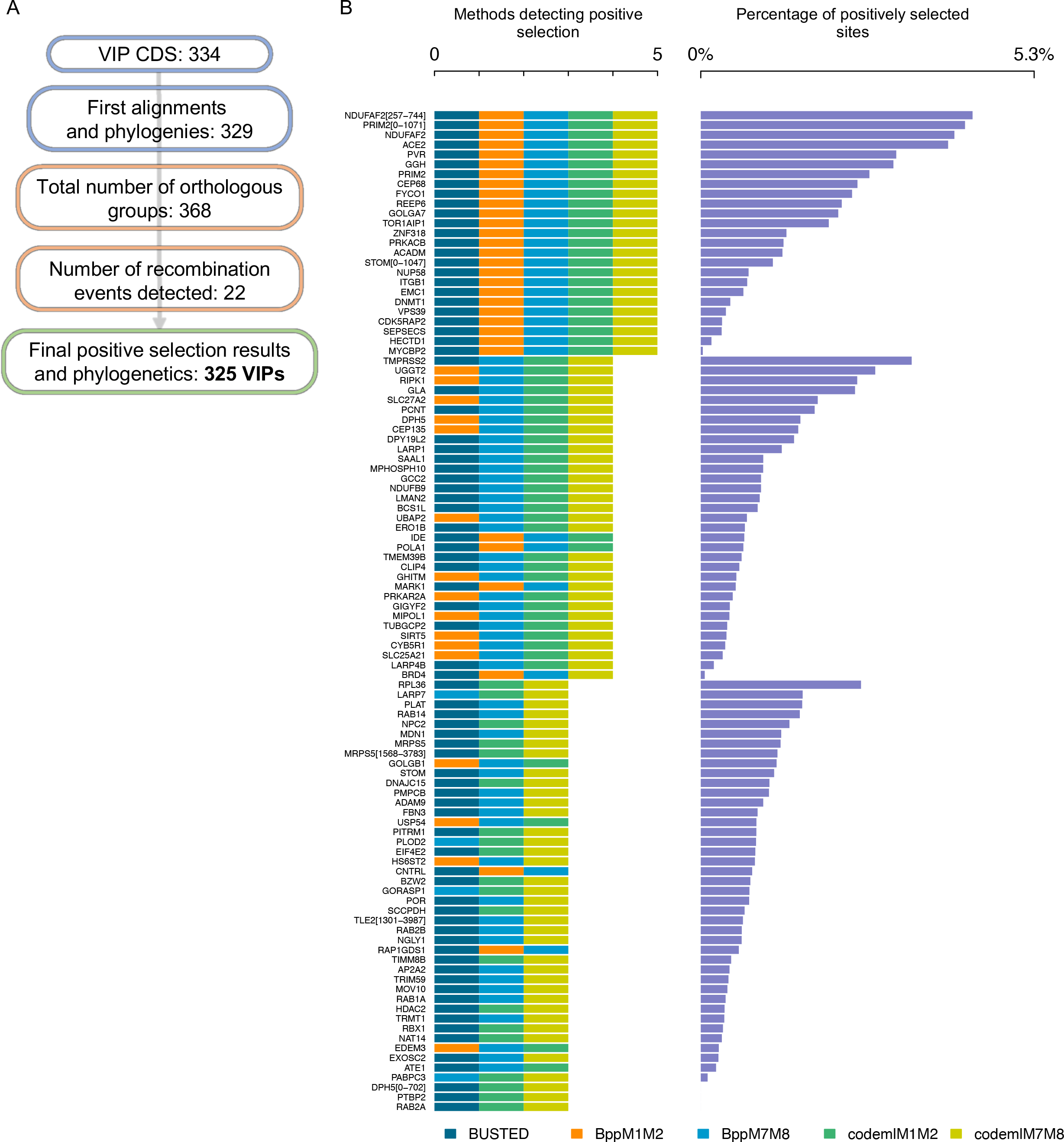
Identification of SARS-CoV-2 interactome with signatures of positive selection in primates. A, Overview of the key steps of the bat VIP DGINN screen workflow. Details in Table S1. B, VIP-encoding genes identified under significant positive selection by at least three methods in DGINN (embedded legend). The percentage of positively selected sites in each VIP is shown on the right panel. Of note, seven genes are false positives due to erroneous sequences or alignments: EMC1 (ER membrane protein complex subunit 1), MOV10 (Mov10 RISC complex RNA helicase), POR (cytochrome p450 oxidoreductase), PITRM1 (pitrilysin metallopeptidase 1), RAB14, RAB2A, and TIMM8B (translocase of inner mitochondrial membrane 8 homolog B).

**Figure S3.**
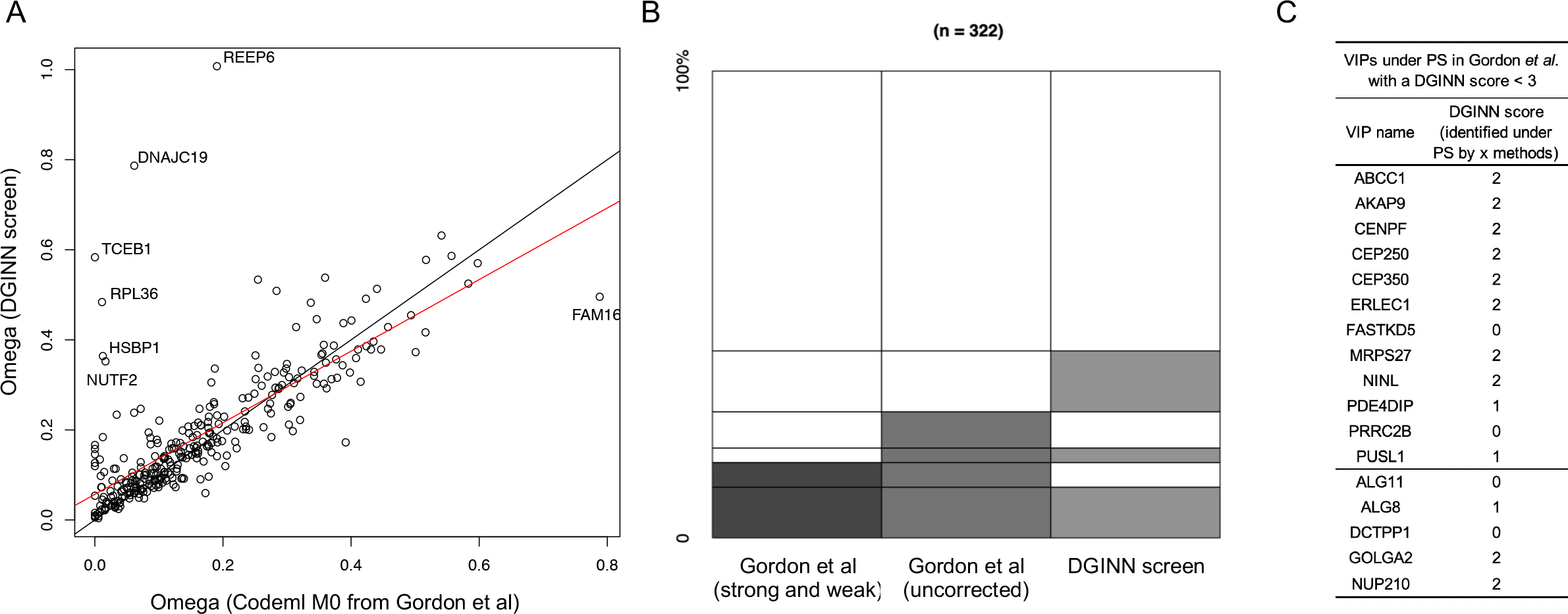
Comparison of primate positive selection analyses between this study and Gordon et al. A, Comparison of the omega (dN/dS) values in PAML Codeml M0 model of the primate VIP genes calculated using the automated DGINN pipeline (y axis) and from Gordon et al study (x axis). Raw data were kindly provided by Janet Young, Fred Hutchinson Cancer Research Center, Seattle, WA, USA. In black, the bisector. In red, the linear regression. B, Comparative analysis of the number of VIPs “under positive selection” in primates. In dark grey, the “strong and weak positively selected genes with benjamini-hochberg correction” from Gordon et al study (Codeml M8 vs M8a, p-value < 0.10); in medium grey, same from “uncorrected p-values”; in light grey, the genes identified by at least three methods in the primate DGINN screen. A total of 322 genes were in common between the two studies. C, VIPs under positive selection in Gordon et al with a DGINN score below three.

**Figure S4.**
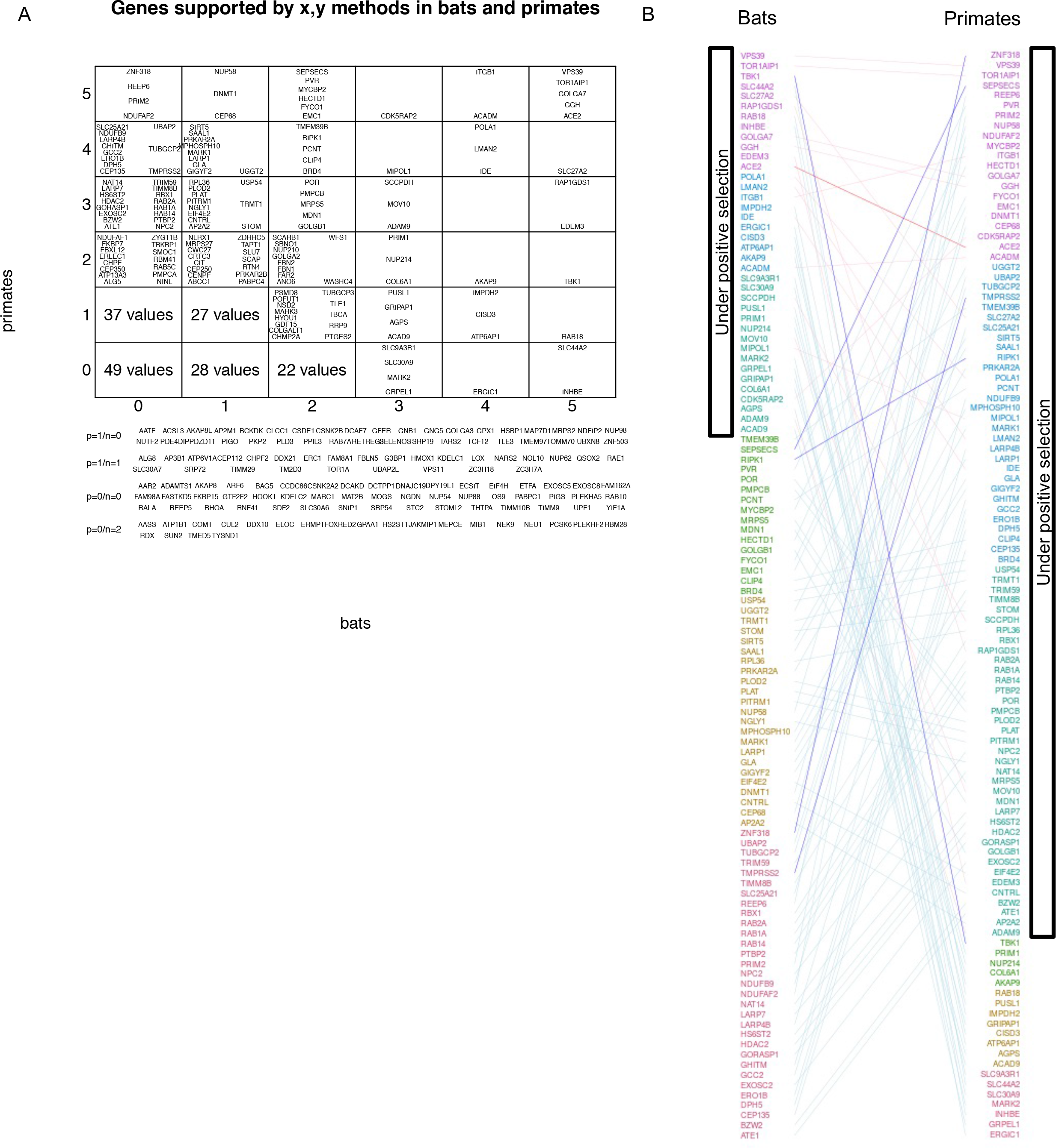
Comparative analyses of adaptive signatures in SARS-CoV-2 interactome in primates and in bats. A, Full table (associated with Figure 1D) showing the genes identified by x,y DGINN methods in bats and primates, respectively. B, Tanglegram of genes under positive selection in bats and primates. At the top, genes with the highest DGINN score (DGINN scores: purple, 5; blue, 4; teal, 3; green, 2; brown, 1; red, 0).

**Figure S5.**
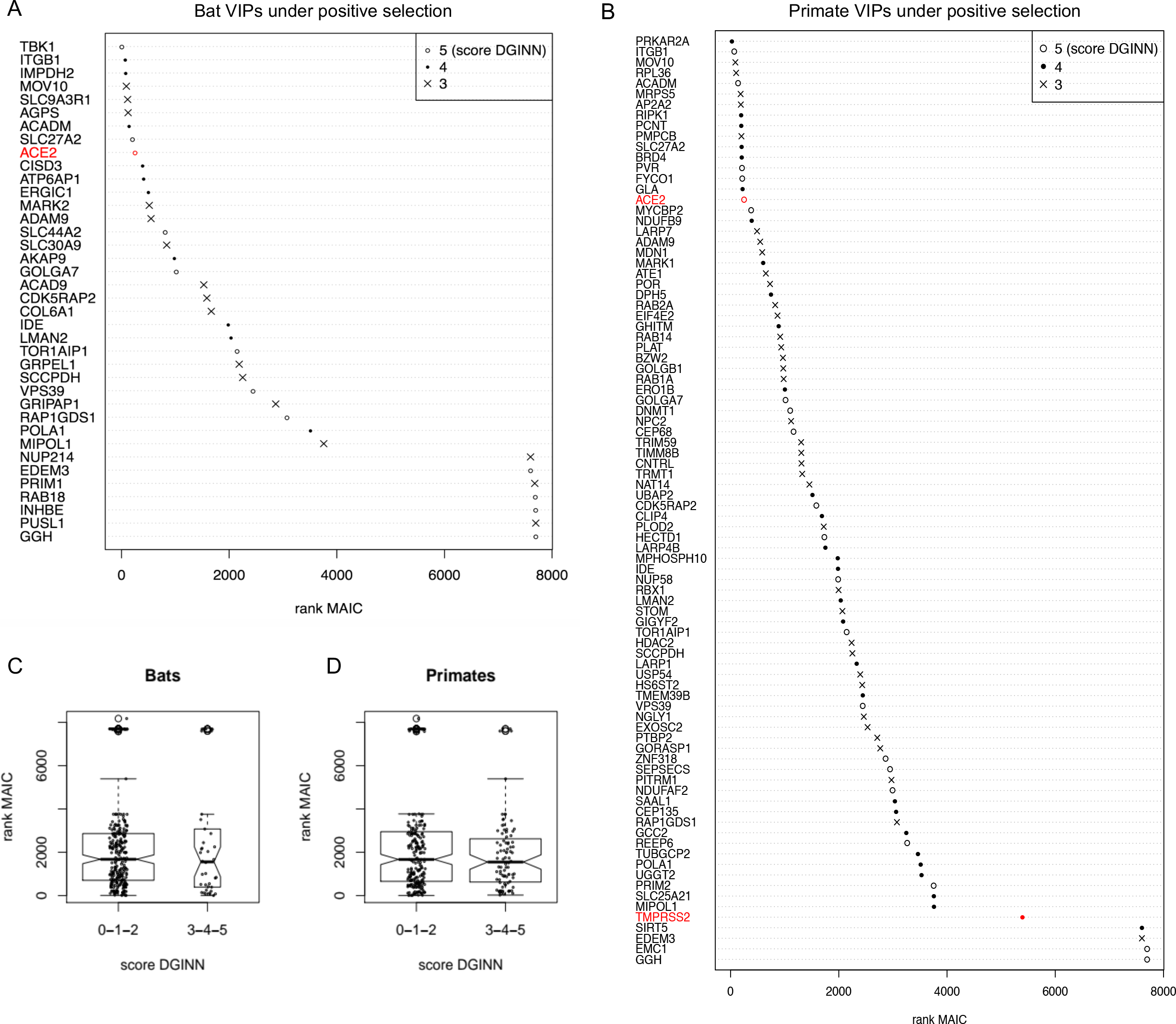
Meta-Analysis by Information Content (MAIC) scores of the VIP genes under positive selection. A-B, MAIC rank of VIPs identified under positive selection (by at least three methods in DGINN; DGINN scores of 3-5) in bats (A) and primates (B). The ACE2 and TMPRSS2 genes are highlighted in red as references. C-D, MAIC rank for all VIPs with, or without, evidence of positive selection (DGINN scores > or = 3, or < 3, respectively) in bats (C) and primates (D).

**Figure S6.**
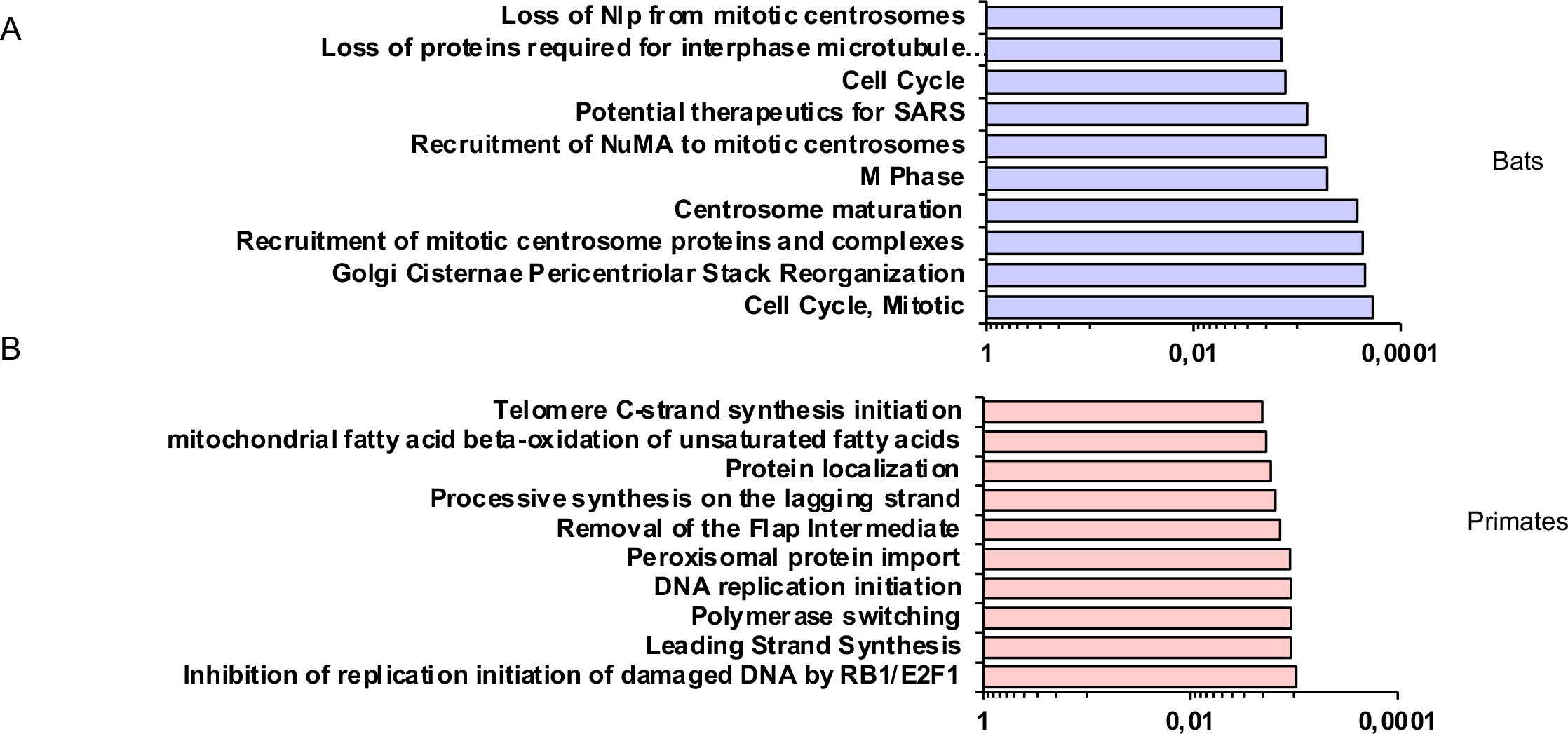
Positively selected VIPs are involved in several biological processes. The graphs present the top 10 biological pathways retrieved after analysis on the Reactome database of bat (A) and primate (B) positively selected VIPs.

**Figure S7.**
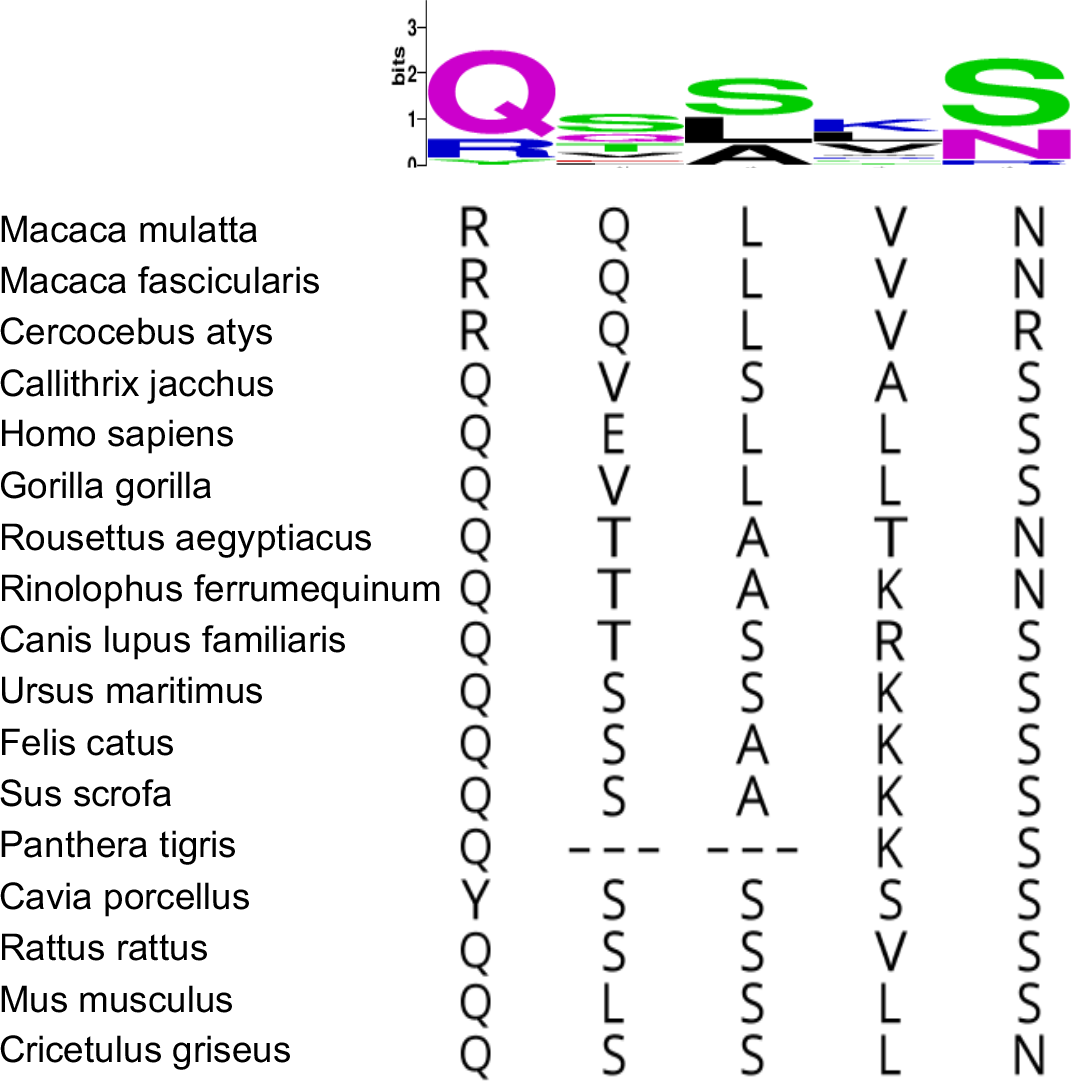
Genetic variation of mammalian TMPRSS2 at the corresponding residues under positive selection in primates. Top, sequence logo of the positively selected sites in mammals that are naturally and experimentally permissive to SARS and MERS coronaviruses. Logos generated using WebLogo based on the alignment of the positively selected sites identified in primates. Bottom, Alignment of the corresponding amino acids in the mammalian species.

**Figure S8.**
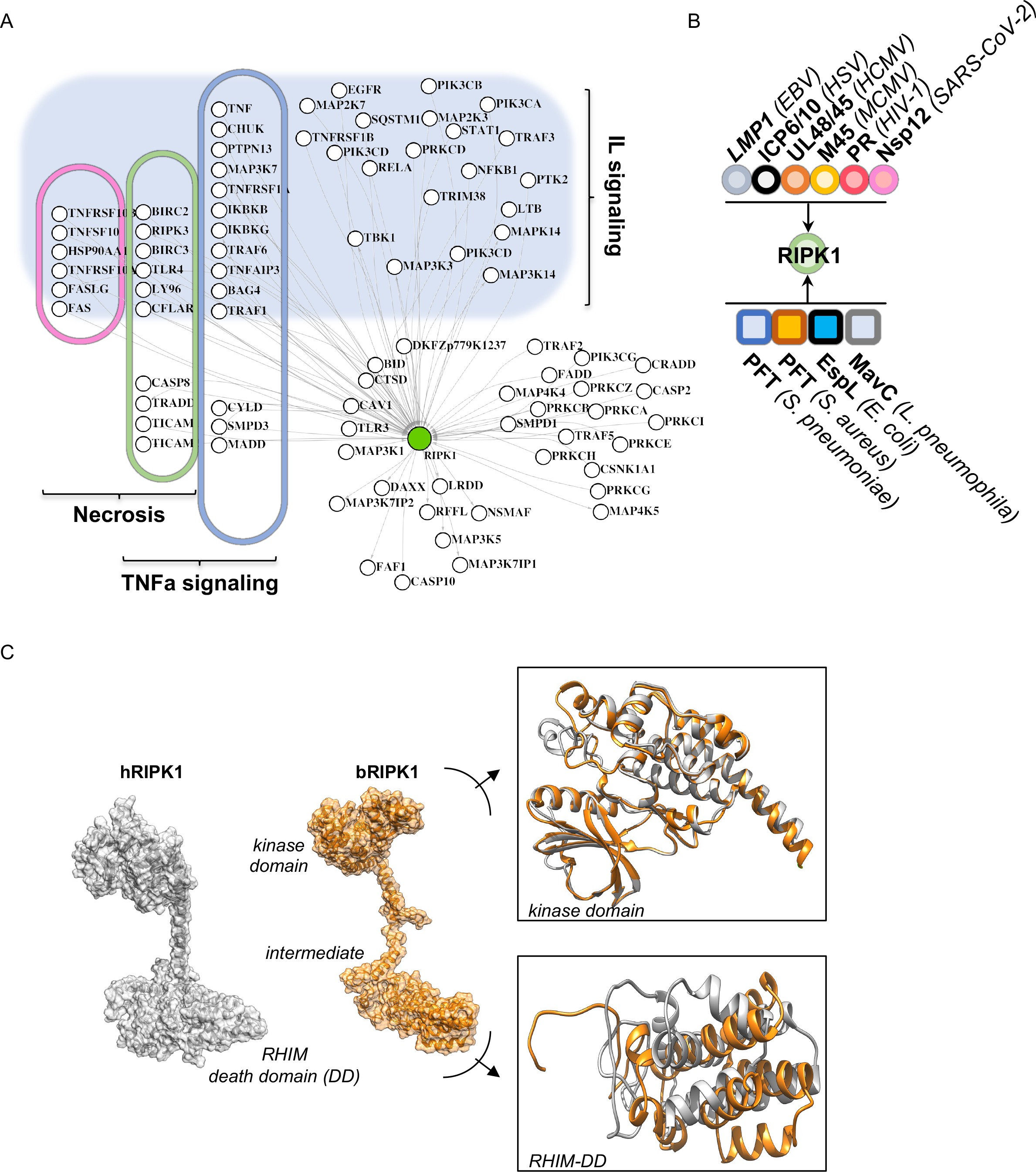
RIPK1 interactome and 3D-RIPK1 structure prediction. A, RIPK1 was used to interrogate the Reactome database and to retrieve RIPK1 cellular interactors that were subsequently subdivided according to their involvement in the indicated pathways. B, Described RIPK1 microbial antagonists or interactors from viruses (top) and bacteria (bottom). C, Human and *Rhinolophus ferrumequinum* RIPK1 sequences were used to generate 3D-structure prediction models on RaptorX (grey and orange, respectively). Magnified views of the structural homologies between the indicated domains.

**Figure S9.**
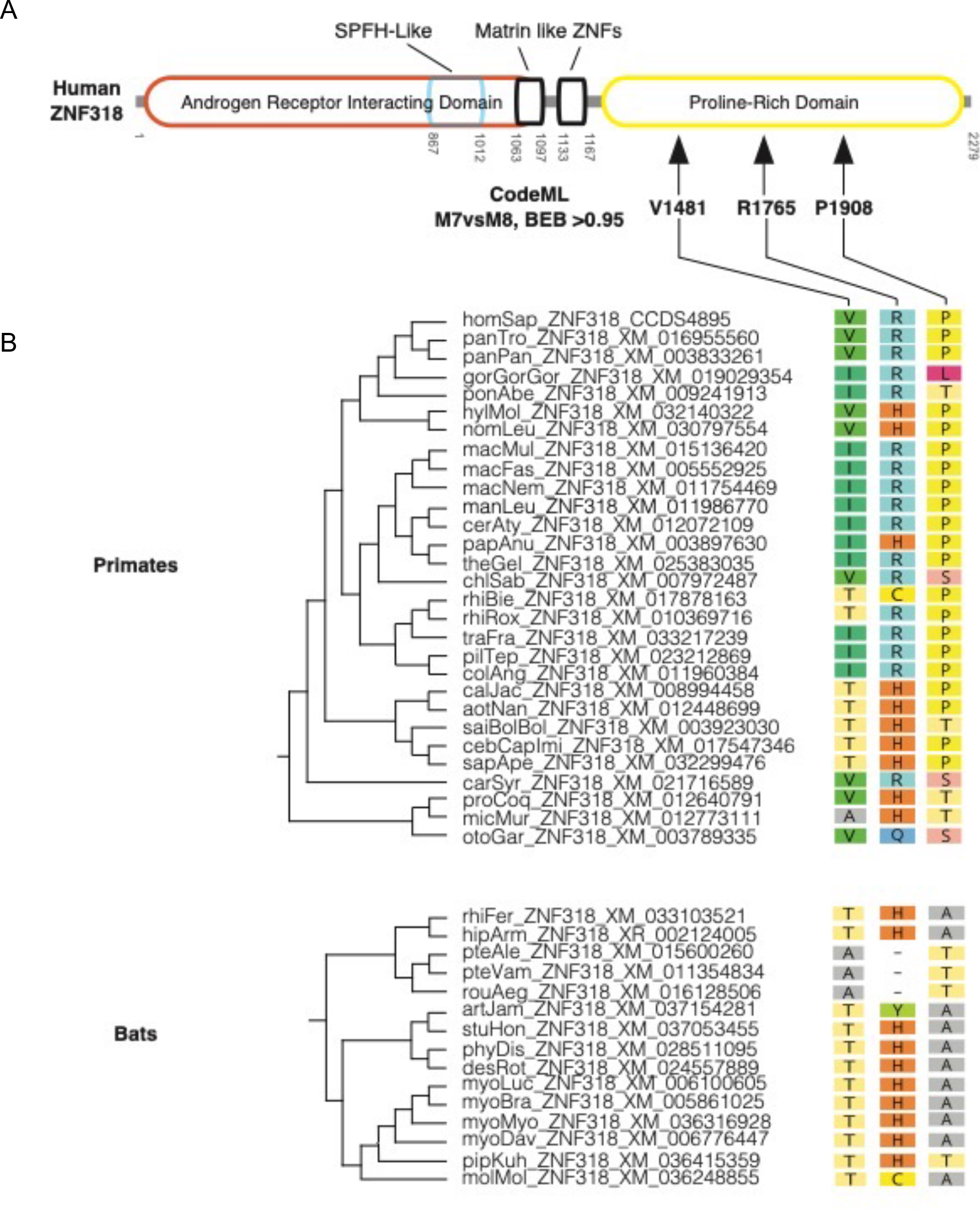
ZNF318 has undergone positive selection at three sites in primates, but not in bats. A, Schematic representation of human ZFN318 protein domains. Numbering of residues is relative to the human full-length protein sequence. The three residues subjected to positive selection in primates are shown with arrows. B, Amino-acids found at rapidly evolving sites across ZFN318 primate orthologs used in this study (top). Orthologs are organized according to the accepted species phylogeny. Bottom, same as top with the corresponding residues in bats. Dashes indicate an alignment gap due to an indel.

**Figure S10.**
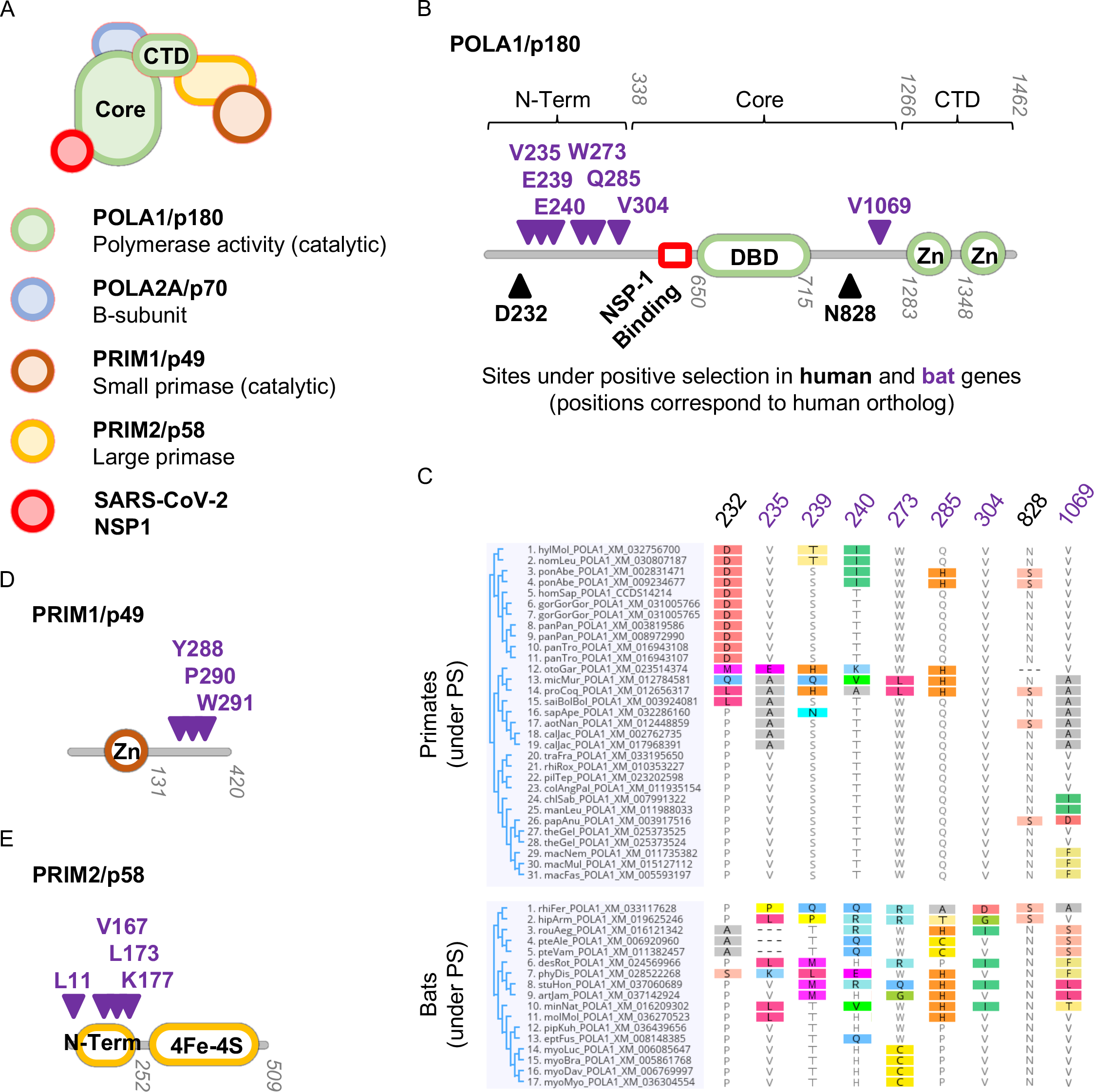
Rapid evolution of the Prim-Pol primase complex (POLA1, PRIM1, PRIM2) in primates and bats. A, Schematic representation of human DNA primase complex. B, D, & E, Diagrams of predicted domains for POLA1, PRIM1, and PRIM2, respectively. Sites under positive selection in primates are represented by black triangles and sites under positive selection in bats are represented by purple triangles (Table 1). Codon numbering based on Homo sapiens genes. C, POLA1 amino acid variation at the positively selected sites in primates (top) and bats (bottom). Codon numbering and coloring as in B.

## Supplementary Tables

**Table S1.** Number of genes screened in the initial primate and bat automatic screens – associated to Figures S1-S2.

|  |  | **Primates** | **Bats** |
| --- | --- | --- | --- |
| **Initial dataset (Gordon et al., 2020 + ACE2 + TMPRSS2)** |  | 334 | 334 |
| **Phylogenetic analyses** | Failed runs | 5 | 4 |
|  | Duplicated genes retrieved by DGINN | 39 | 5 |
|  | Complete alignments and phylogenies | 368 | 335 |
| **Positive selection analyses** | Results from 0 to 4 (out of 5) DGINN methods | 43 | 11 |
|  | Complete results (5/5 PS methods) | 325 | 324 |

**Table S2.** SARS-CoV-2 interacting proteins with no other known virus interactors

|  |
| --- |
| HGNC names of the SARS-COV-2 VIPs with no other known virus interactors (*, except with other coronaviruses) |
| **ACE2\*** |
| GOLGA7* |
| **PRIM1\*** |
| EIF4E2* |
| GORASP1* |
| MARK1* |
| TRIM59* |
| CISD3 |
| GGH |
| INHBE |
| PUSL1 |
| DPH5 |
| GCC2 |
| HS6ST2 |
| NUP58 |
| PLAT |
| PTBP2 |
| TMEM39B |
| **TMPRSS2\*** |

